# Cognitive Graphs of Latent Structure in Rostral Anterior Cingulate Cortex

**DOI:** 10.1101/2025.04.18.649507

**Authors:** Maksym Manakov, Mikhail Proskurin, Hanqing Wang, Elena Kuleshova, Andy Lustig, Reza Behnam, Shaul Druckmann, D. Gowanlock R. Tervo, Alla Y. Karpova

## Abstract

Mental maps of environmental structure enable the flexibility that defines intelligent behavior. We describe a striking internal representation in rat prefrontal cortex that exhibits hallmarks of a workspace for constructing goal-specific, cognitive graphs for sequences of abstract states that animals must traverse through their actions. As rats uncover — unguided — complex sequential patterns, neural ensemble activity in rostral Anterior Cingulate Cortex develops a highly structured yet compact, scalable representation anchored in states at the start and end of self-organized sequences. Graphs are organized to reflect relational similarities across contexts, and individually, permit flexible refinement of component states. The representation’s crystalline organization permits instantaneous inference of the animal’s current state within self-generated sequences, offering insights into the algorithmic and representational principles underlying unguided parsing of open-ended problems.

## Main Text

A hallmark of biological intelligence is the ability to achieve diverse goals in uncertain and dynamic environments. Such intellectual versatility allows humans to adapt their finger movements to different smartphone apps, and animals to customize foraging paths based on the estimated distribution of resources, predators and competition. This aspect of biological intelligence is underpinned by the brain’s capacity to develop a causal and generalizable understanding of the environment by discerning the often-latent relational structure of experienced events (*1–3*), representing that structure as a mental model (*4*), and adaptively using it to achieve specific outcomes (*5*). In the mammalian brain, frontal cortical areas (*6–9*) and the hippocampal formation (*10, 11*) have been most strongly implicated in the construction and maintenance of mental models that organize accumulated knowledge into relational structures in service of behavioral flexibility, often referred to as cognitive maps (*5, 10, 12, 13*). Exemplified most strikingly in the spatial domain by place cells in the hippocampus (*14*) and grid cells in the entorhinal cortex (*15*), cognitive maps are thought to support mental exploration through chains of instantiated ‘states’ (*16*) to form robust predictions, plan future routes and discover shortcuts, as well as to make rapid inferences in structurally related settings. While the concept of a map is most intuitive and literal for spatial problems in Euclidean space, its graph-like generalization — with states as nodes linked by edges — is also suitable for reasoning about diverse abstract entities, both spatial and non-spatial (e.g., sequential steps that lead to a goal, the relationship between individuals within a social network, etc.) (*17, 18*).

The discovery of place and grid cells has provided the conceptual and experimental traction necessary to uncover broad principles of knowledge organization within cognitive maps (*10, 19–22*). This contrasts with our comparatively impoverished understanding of the relevant neural representations within frontal brain regions, where numerous physiological studies have revealed complex, mixed selectivity for a multitude of experimental parameters (*23, 24*). Recent theoretical accounts have argued that frontal regions may be particularly important for creating a map focused selectively on states relevant to the animal’s current objective (*25, 26*). While a wealth of experimental findings documents the encoding of rich information relevant to acquiring knowledge relevant to a specific objective and structuring behavior accordingly (*27–38*), we have yet to discover clear representational principles in frontal cortex — analogous to the crystalline representations within the hippocampal formation — that would further elucidate component computations in cognitive mapping.

Some insights into the nature of representations within frontal cortex derive from neuropsychological accounts of deficits in patients with anterior prefrontal cortical lesions. These patients can perform familiar tasks, and can learn new tasks comprising sequences of pre-specified subgoals if they are provided with detailed step-by-step instruction, but they struggle to reason, unguided, through multi-step open-ended problems that lack a prespecified set of possible solutions (*39–41*). Thus, a core possible deficit following anterior frontal lesions includes an inability to distill, out of an overwhelming amount of information about the world, a small set of relevant causes (or states) and a concise cognitive graph that relates those states and actions that could be sequenced to traverse them to achieve a particular end goal. To investigate whether such graphs are indeed represented in the frontal cortex, we developed a task in rats that requires an entirely unguided discovery of latent sequential patterns while monitoring population activity in the Anterior Cingulate Cortex to reveal a striking form of representation that exhibits hallmarks of a cognitive graph ‘workspace’ useful for parsing large, open-ended settings.

### Unguided latent sequence discovery framework

We first designed a behavioral paradigm that offered rats no explicit guidance, predetermined solutions or step-by-step feedback, thereby encouraging animals to extract relevant latent structure and rely on it for their actions. Rats learned to discover, unguided, through serendipitous reinforcement that a sequence of compound actions — symbolized by ‘L’ or ‘R’ — could lead to rewards (*42*). Reinforcement was made available whenever an experimenter-determined target pattern (e.g., ‘RRRL’) was spontaneously produced by the rat. To engender a symbolic sequence representation, we defined ‘L’ and ‘R’ as paired center-to-side port transitions in a simple chamber with three nose ports arranged left to right (Fig. 1A). Requiring an initiating center port entry for every symbol ensured that identical symbols (e.g., two ‘R’s) were produced with relatively consistent movement patterns rather than varying based on prior position. This setup prevented confounding motor strategies for sequence generation and ensured that the rats had to independently discover both how to generate individual symbols and how to reconstruct entire target sequences from reinforcement alone (Fig. 1B). Crucially, because the length of the latent target pattern could vary, there were no predetermined solutions, and the only feedback came after a behavioral sequence matched the target pattern, thereby ensuring that animals had to uncover the latent pattern through self-directed exploration.

**Figure 1.**
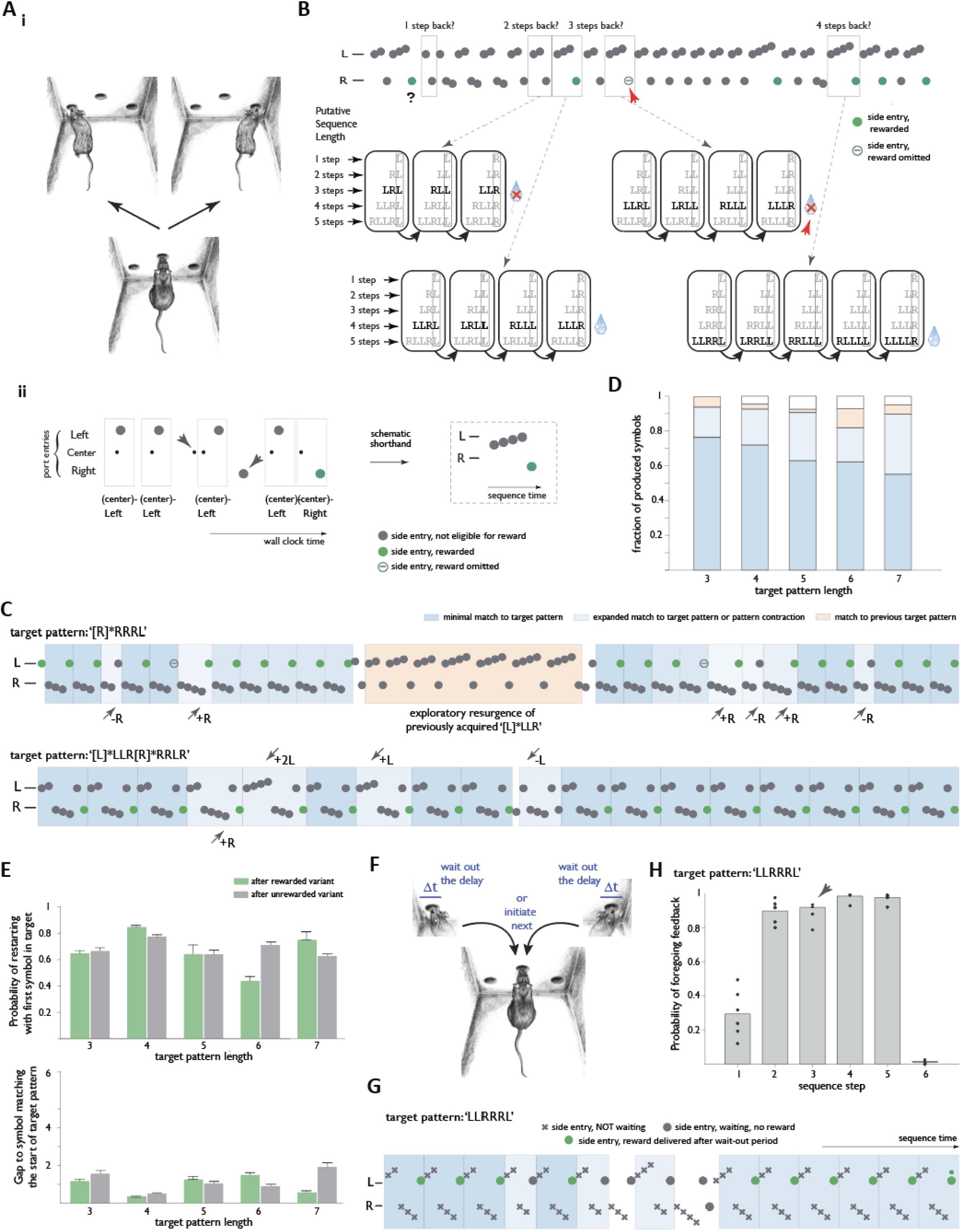
Unguided latent sequential pattern discovery framework prompts rats to develop cognitive sequence abstractions adaptively aligned with target patterns. (**A**) Concept. i, ‘L’ and ‘R’ are defined as paired center-to-side port transitions in a simple chamber with three nose ports ii, The animal is eligible to receive a reward only if an uncued experimenter-defined target pattern (here, ‘LLLR’) appears in the symbol stream produced by the animal. The behavioral stream is schematized by the sequence of L and R port entries, with rewarded entries, reflecting termination of the target sequence, indicated in green. For simplicity, behavioral data is presented throughout the rest of the paper in sequence time with center port entries omitted. (**B**). Illustration of the complexity of mapping serendipitously acquired rewards to useful state representations in the absence of cues that indicate sequence starts and ends. Neither the length nor the composition of the target pattern is indicated, and rewards are occasionally omitted, necessitating discovery of the relevant target sequences through exploration. Here, for example, a reward (green circle marked “?”) is encountered after the rat produces the behavioral stream “…LLRLLLR”, but how far back the target starts before the terminal R is not signaled; the various exploratory sequences embedded within the subsequent behavioral stream (boxed patterns) gradually inform and constrain how far back the target sequence starts. (**C**) Prevalence, in two example behavioral traces from the same animal, of sequence variants that either represent minimal matches to the target pattern (darker blue), expanded matches and pattern contractions (lighter blue), or previously acquired target patterns (light orange). (**D**) Prevalence of the above three sequence classes across the behavioral dataset for target patterns of different lengths [n= 15,14,11,9 and 14 sessions; N=5,5,4,3 and 4 animals for target patterns of length 3-7 respectively]. (**E**) Probability of (top panel) and gap to (bottom panel) restarting at the correct starting symbol of the target pattern following rewarded and unrewarded sequence variants. (**F**) Delayed feedback concept. Reward on eligible trials is delivered only if the animal waits in the side port for the duration of a multi-second delay. (**G**) Example behavioral trace for an animal that has acquired ‘LLRRRL’ pattern and was challenged with delayed reward. (**H**). Probability that animals forego feedback on individual steps of all ‘LLRRRL’s in symbol stream. Note consistent pausing at the end of the length=6 target, and lack of pausing at other steps including the junction between ‘LLR’ and ‘RRL’ (arrowhead) [n=6 sessions, N=3 animals].

Several features of this design made it well-suited for studying the generation of abstract sequences in pursuit of specific objectives. With symbol identity abstractable away from physical locations, sensory cues, or motor patterns, an abstract representation would need only to include the symbolic components of sequences. The small set of symbols increases the likelihood of rats serendipitously matching target patterns even in the absence of any explicit guidance, allowing them to refine their behavior based on reinforcement. Once rewarded, animals can, in principle, work backward to determine how they arrived at the reinforced sequence. This would require that they both maintain a ‘memory buffer’ of past choices and can trace steps from potential start states to the goal state. Despite the absence of external cues to indicate starts and ends, animals can then systematically test and modify (“edit”) their mental representation, potentially a cognitive graph, by shifting their ‘sequence start register’, iteratively refining their hypothesis (Fig. 1B). Changing the target pattern periodically permits experimentally ascertaining the graph’s specificity for a specific objective.

Consistent with our prior studies (*42*), we found that rats readily discovered target patterns of length 3, such as ‘[L]*LLR’ and ‘[R]*RRL’, where ‘[L]*’ and ‘[R]*’ indicate that the corresponding symbol could be repeated beyond the minimal pattern. To further challenge the animals, we increased latent structure complexity by lengthening the target patterns or concatenating previously discovered patterns to construct longer ones. For example, Fig. 1C displays the behavior of a rat that discovered the patterns ‘[R]*RRRL’ and ‘[L]*LLR[R]*RRRLR’ that were latent targets in different sessions. These patterns represent an extension of ‘[R]*RRL’ and a concatenation of ‘[L]*LLR’ and ‘[R]*RRLR’, respectively. More broadly, even in this open-ended setting, where target patterns ranged from 3 to 7 symbols in length, most (>90%) of the rats’ behavioral sequences reflected a structured relationship to the target patterns. Their behavior included minimal matches to the target (e.g., producing ‘RRRL’ when ‘[R]*RRRL’ was the target pattern), expansions (e.g. ‘**R**RRRL’), contractions (e.g. ‘RRL’), and reuse of previously discovered patterns (Fig. 1C,D; Materials and methods). Additionally, rats typically reinitiated their sequences with the appropriate starting symbol of the target pattern, regardless of whether they had just completed a rewarded match, an unrewarded contraction, or even a match for which reward was intentionally omitted (Fig. 1E). This behavior suggests that rats were not solely reliant on reinforcement to recognize when they had reached the goal state. Together, these findings indicate that the rats structured their behavior at the level of cognitive sequences composed of abstract ‘L’ and ‘R’ symbols, rather than merely producing ‘L’ and ‘R’ choices with some fixed frequency.

We next sought to directly determine whether rats impose mental punctuation on their otherwise continuous stream of ‘L’ and ‘R’ symbols to define sequence ‘chunks’ (*43–46*) that they can flexibly adjust to match different target patterns. Intuitively, an animal with a structured mental representation of a target pattern should expect a reward only upon reaching the mentally defined endpoint of the pattern match, but not in the middle. Likewise, failure to receive a reward should be informative only after reaching the endpoint. For example, if a rat learns that ‘LLR’ and ‘RRL’ are target patterns, it should punctuate them as ‘LLR●’ and ‘RRL●’, treating each as a distinct chunk. However, if the rat later discovers that ‘LLRRRL’ is a useful pattern, it should update its mental model, punctuating the sequence as ‘LLRRRL●’ rather than ‘LLR●RRL●’. Such an ability to flexibly restructure the boundaries of sequence chunks would suggest that rats dynamically adjust their cognitive representations to align with newly reinforced patterns.

How might we directly read out an animal’s mental punctuation? We reasoned that animals would aim to complete the steps of a particular sequence chunk as quickly as possible and modified the paradigm by requiring a pause to receive reward/no-reward feedback (*47, 48*). Specifically, we introduced a substantial delay (typically fixed at 1-3 sec) between a side port entry and the onset of reward delivery for terminal steps completing a target pattern match (Fig. 1F). Since expert rats typically executed individual steps in under a second, this delay created an incentive to skip feedback on intermediate steps within indivisible chunks. After some experience with this paradigm, rats that had previously discovered ‘RRL’ and ‘LLR’ patterns displayed a robust preferential pausing after the third step, explicitly punctuating their symbol streams as ‘RRL●’s and ‘LLR ●’s (fig. S1A). We then challenged a subset of these animals to discover the more complex ‘LLRRRL’ pattern, this time without imposing feedback delays during the search phase. Once their symbol streams contained ‘LLRRRL’ matches above chance levels (i.e., exceeding random recombination of the three-step components, fig. S1B), we reintroduced feedback delays, revealing that rats now explicitly punctuated sequences as ‘LLRRRL●’ (Fig.1G-H). This result demonstrated that rats flexibly adjusted their mental punctuation to align with the newly acquired complex target pattern.

In summary, before carrying out neurophysiological investigations of prefrontal representations, we trained rats to develop, through self-guided exploration, a series of sequence abstractions consisting of structured ‘L’ and ‘R’ patterns. By systematically varying target patterns—both within and across rats—in terms of length, complexity, and construction rules (i.e., via lengthening or concatenation), we established that our rats adaptively re-structured their representations accordingly. This setting thus created an opportunity to probe both the overall nature of these representations and any features that might have facilitated the rats’ discovery of pattern extensions and concatenations.

### Graph-like neural representations of sequence states in Rostral Anterior Cingulate Cortex, but not Ventromedial Prefrontal Cortex

Motivated by our behavioral evidence for abstract sequence representations and reasoning that these would be in the form of neural population activity (*34, 46, 49–51*) in the rostral prefrontal cortex (*52–56*), we next sought to record from large ensembles of neurons in this region in rats. Specifically, we recorded simultaneously from hundreds of neurons using high-density Neuropixels probes in rostral anterior cingulate cortex (rACC; area 32d, Fig. 2A) while animals repeatedly adjusted to unsignaled changes in target pattern structure. To minimize confounds from movement-related activity as animals made left or right choices, we analyzed activity when the animal was at the center port that initiated every ‘L’ and ‘R’ step in the behavioral stream. This provided a consistent temporal reference point to compare neural dynamics across different steps during sequence production. We began by examining the structure of population activity within a high-dimensional neural space, where each dimension corresponds to the spike count of a single neuron during a fixed 500 ms window (Fig. 2B; window center aligned on port exit due to different port dwell times across animals, see Materials and Methods). In this space, neural population activity at each moment is captured by a single point. Given rats’ continuous and varied movements over several hours, no specific structure in these activity patterns across a session would be expected by default.

**Figure 2.**
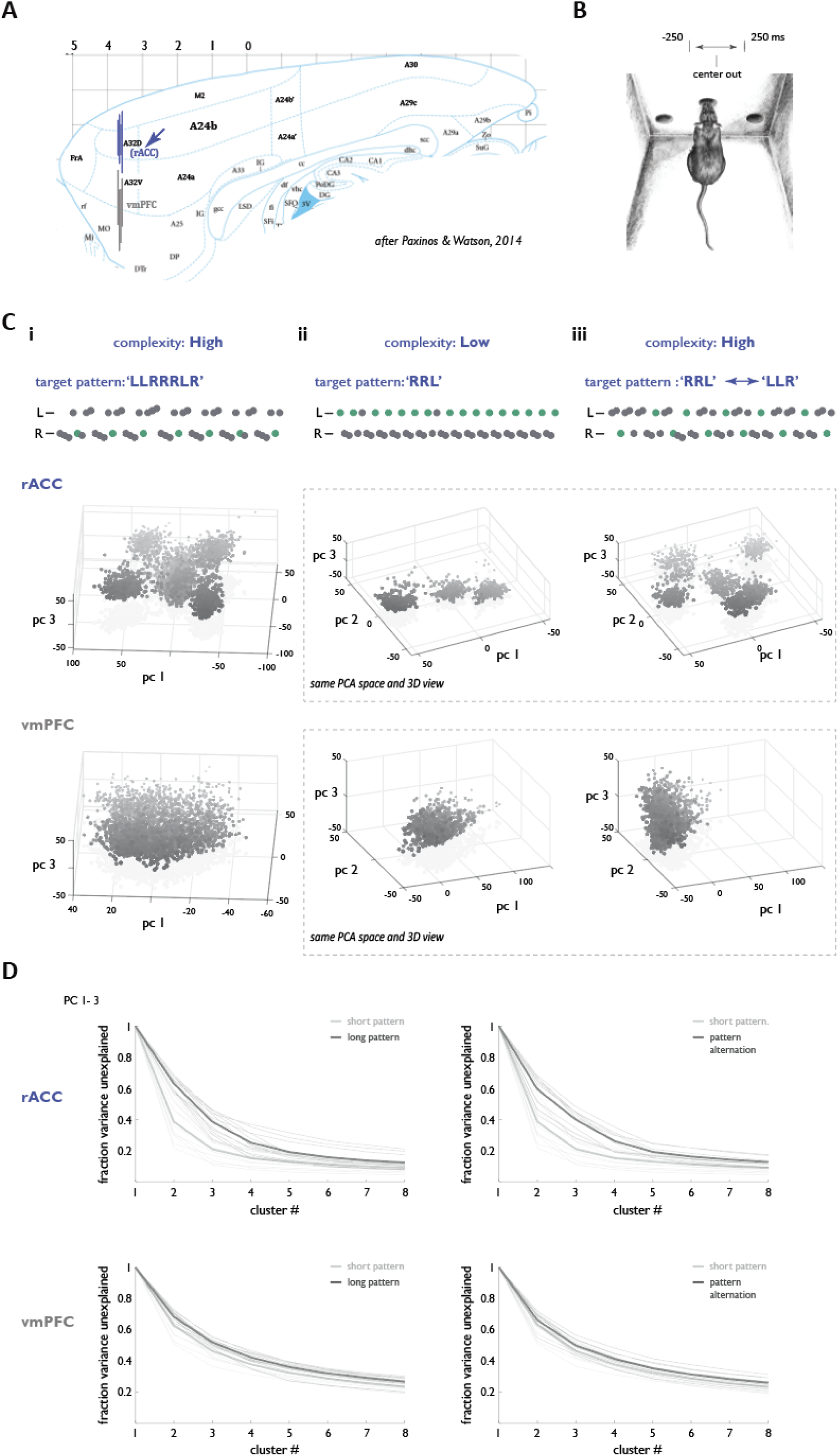
A form of neural punctuation engenders a highly structured rACC representation that scales with latent structure complexity. (**A**) Approximate location of rACC (blue) and vmPFC (grey) recording sites. (**B**) Schematic of analysis window alignment. (**C**) Reference behavioral traces (top panel), population activity plotted in the space of the first three principal components for rACC (middle panel) and vmPFC (bottom panel) for an example session where latent structure was a long sequence (i), a short sequence (ii), or alternation between two short sequences (iii). Note that each point corresponds to population activity at the center port of a single sequence step. Note also that shading is used solely to facilitate visualization of depth. (**D**) Fraction of rACC (top row) or vmPFC (bottom row) data variance left unexplained by a model with different cluster numbers for sessions with long vs short sequences as latent targets (left row), [n=12 sessions, N-=4 animals], and session with blocks of a single short sequence or forced alternation between two short sequences as latent targets (right row), [n=9 sessions, N=3 animals]. Note that sample sizes were matched for comparisons within the same session. Thin lines: individual sessions. Thick lines: session averages.

To visualize this high-dimensional activity while preserving its geometry, we applied principal component analysis (PCA), reducing the data to three dimensions that captured the most variance. Strikingly, rACC ensemble activity was organized into a small number of discrete, well-separated clusters in PC space (Fig. 2C, top middle panel, Movie S1). In sessions where rats discovered latent target patterns of order 3, a simple three-cluster model explained 79.1 ± 1.9 % of the variance in the space of the first 3 PCs and 59.2 ± 4.2 % of the variance in the high-dimensional space (defined here as the subspace capturing 75% of total activity variance; see Materials and Methods). In contrast, recordings from ventromedial prefrontal cortex (vmPFC) — deeper along the medial wall — showed no such clustering, despite sampling an equivalent number of neurons. In vmPFC, the same model explained only 53.6 +/− 1.5 % of variance in the space of the first 3 PCs and 27.8 +/− 1.1 % in the high-dimensional space (Fig. 2C, bottom middle panel, Movie S2). These results suggest that clustered neural representations are not a trivial feature of prefrontal circuits, nor are they guaranteed simply by the production of sequences. Importantly, rACC clustering was equally strong when population activity was aligned to side port entry (variance explained by a three-cluster model: 78.5 ± 1.8%, 58.3 +/− 3.1%), indicating that the representational structure was not restricted to a single moment within the behavioral sequence. These findings suggest that the rACC portion of the medial prefrontal cortex may selectively encode a compact, graph-like state representation of animals’ self-discovered sequence patterns, reflecting an internal model of the latent structure guiding behavior. This interpretation leads to three key predictions: that the graph-like representation should scale with complexity (i.e., more ‘nodes’ or clusters for more complex latent target patterns), be highly context-specific (i.e. new graphs constructed whenever an animal perceives the target pattern to have changed), and mirror the self-organized structure of the animal’s behavior.

To assess — in an unsupervised fashion — the number of distinct nodes represented in rACC activity across contexts of different target pattern complexity, we fit Gaussian mixture models to neural population activity in each session and quantified the variance left unexplained as a function of different cluster numbers of clusters (see Materials and Methods). Consistent with the prediction of scaling with target pattern complexity, more clusters were required to explain the neural data to the same degree of accuracy in sessions where animals had discovered longer patterns (Fig. 2C, top left panel, Movie S3). This held true whether the analysis was done using the 3 first PC dimensions (Fig. 2D, 61.4+/−1.9% of data variance in high complexity setting explained by a model with 3 clusters compared with 79.1+/−1.9% in low complexity setting, p< 10^-4^, Wilcoxon rank-sum test) or in the high-dimensional space (fig. S2, 40.0+/−3.4% for high complexity vs 59.2+/−4.2% for low complexity, p<0.005, Wilcoxon rank-sum test,, n=23 sessions, N=4 animals). One caveat is that these comparisons typically involved sessions separated by days or even weeks, and thus potentially sampled from slightly different neural populations. To directly compare representations from the same neurons, we designed a within-session manipulation of latent structure complexity via a ‘forced pattern alternation’ paradigm. Instead of lengthening the target pattern, we required animals in this paradigm to alternate between two known patterns (‘RRL’ and ‘LLR’), such that after matching one and being rewarded, they had to switch to match the other before being rewarded again (Fig. 2C, top right panel, Movies S4; Materials and Methods). This manipulation increased latent structure complexity without requiring animals to search for longer target patterns. As expected, variance-left-unexplained curves again shifted toward requiring more clusters, indicating an increased number of distinguishable states (Fig. 2D; e.g., 60.0 ± 2.4% of variance explained in high complexity vs. 79.0 ± 1.9% in low complexity using 3-PC space, p < 10⁻^3^, Wilcoxon rank-sum test; fig. S2 for high-dimensional results, n = 9 sessions, N =3 animals). These findings further support the idea that rACC representations scale flexibly with demands imposed by the complexity of environment’s latent structure.

We next investigated whether rACC representations are sensitive to behavioral context, specifically whether distinct discovered latent patterns (‘RRL’ vs. ‘LLR’) are encoded in separate neural graphs. In these sessions, rats adjusted to repeated unsignaled switches between ‘RRL’ and ‘LLR’ blocks (Fig. 3A). If the rACC tracks context explicitly, then a combined dataset containing sequences from both contexts should require more clusters to explain the same amount of variance as a dataset from a single context, even when total behavioral symbol count is held constant. Consistent with this prediction, we found that significantly more clusters were needed to explain activity when symbol streams from both ‘RRL’ and ‘LLR’ contexts were pooled, compared to symbol streams from a single ‘LLR’ context (Fig. 3B, Movie S5; fig. S3). This suggests that the rACC instantiates more distinct nodes when representing two contexts, even though the corresponding sequences comprise the same basic symbols (‘L’ and ‘R’). Further, we found that activity nodes were not reused across contexts — each behavioral context was associated with a distinct set of clusters, even though the symbol-level actions were identical (4.33+/− 0.79% of activity patterns corresponding to component symbols in ‘LLR’/’RRL’ pattern matches mapped to the cluster set of the other context). Thus, the rACC cognitive representation is not only structured and scalable, but also tightly linked to changes in the latent structure of the environment.

**Figure 3.**
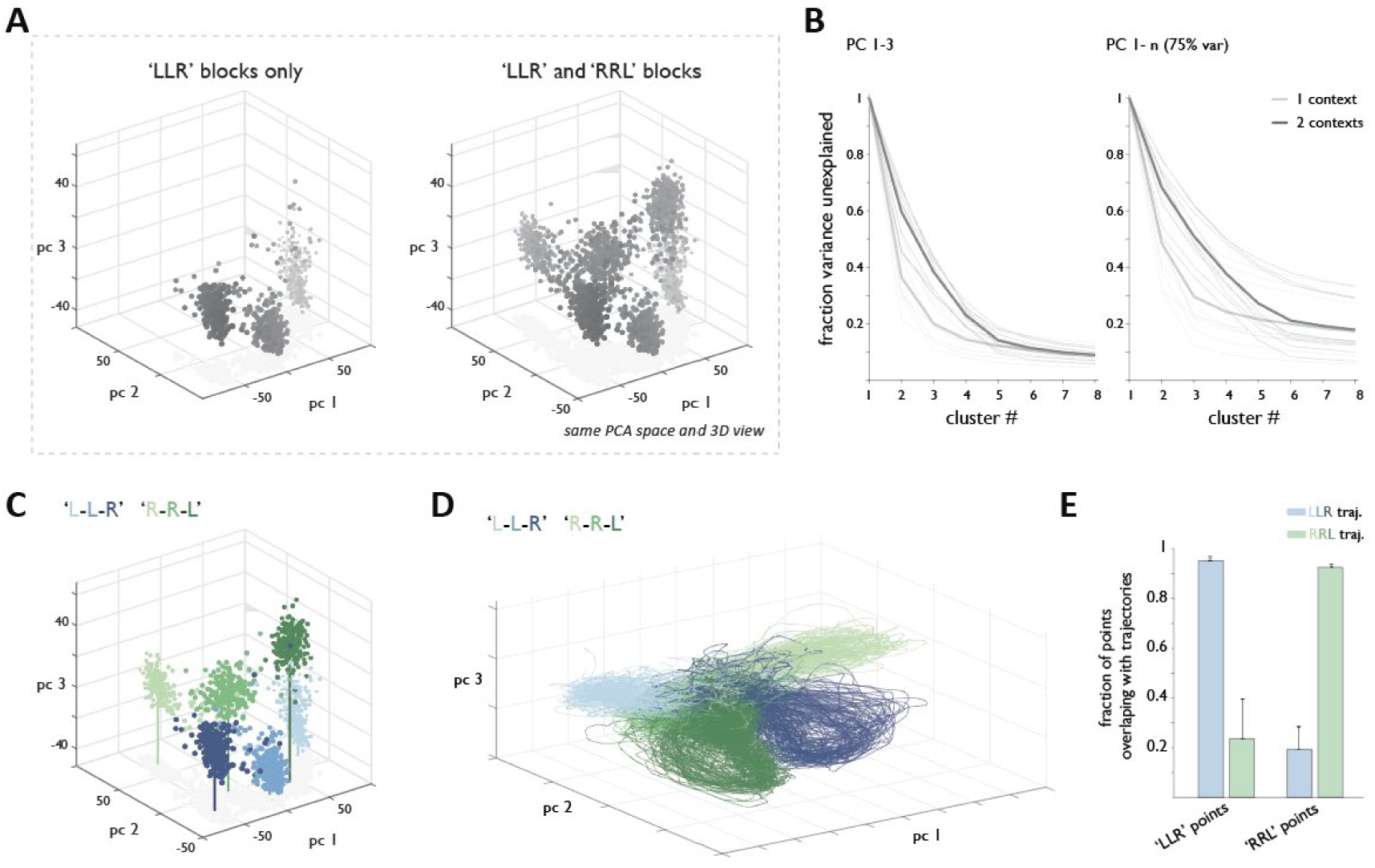
rACC representation is context specific. (**A**) Neural population geometry for ‘LLR’ blocks only (left panel), or ‘LLR’ blocks and ‘RRL’ blocks combined (right panel), plotted in the space of the first three principal components for the entire session. (**B**) Fraction of data variance left unexplained by a model with different cluster numbers for data taken either just from ‘LLR’ blocks or from both ‘LLR’ and ‘RRL’ blocks, down-sampled to match in sample size. Light grey: ‘LLR’ blocks only, dark grey: ‘LLR’ and ‘RRL’ blocks. Thin lines: individual sessions. Thick lines: session averages. n=9 sessions, N=3 animals. (**C**) Neural geometry labeled by block type and sequence step. (**D**) Full sequence trajectories (excluding reward collection) in the same PCA space as (C). (**E**) Fraction of points (activity patterns) in ‘LLR’ and ‘RRL’ contexts overlapping with the space occupied by population activity trajectories during production of ‘LLR’ (blue bars) and ‘RRL’ (green bars) target pattern matches.

We also examined in greater detail how individual activity states relate to behavior separately within ‘RRL’ and ‘LLR’ contexts. Across different iterations of target pattern execution, ensemble activity corresponding to a given step of the sequence tended to fall within the same cluster, indicating consistent representations for each step within a sequence (Fig. 3C, Movie S6, closest cluster matched for 95.6% of points corresponding to individual steps across different sequence instances). We next considered the full trajectory of neural population activity during sequence execution, assessing any possible overlap by defining a per-context region in the space of the first three principal components and then analyzing the number of points from the other context within that region (see Materials and Methods). We found that neural trajectories rarely overlapped between behavioral context (Fig. 3D,E, 19.3+/− 9.4% of ‘RRL’ activity patterns mapped to ‘LLR’ trajectory subspace, 23.6+/−15.9% of ‘LLR’ activity patterns mapped to ‘RRL’ trajectory subspace, n=9 sessions, N=3 animals), supporting conclusions from our earlier fixed-window analyses. Combined, the compactness of rACC representations, their scaling with latent structure complexity, contextual specificity, and close correspondence with steps within unguided sequence production argue that nodes in the rACC neural graphs represent key states within animals’ cognitive graphs of latent sequence structure. Moreover, the crystalline nature of the representation opens the door to interpreting the identity of individual behavioral steps in relation to specific neural graph nodes for instantaneous state inference. Doing so in more complex settings and during learning, however, would require a generalized framework for mapping individual activity nodes onto the topology of expanding cognitive state spaces. This raises the question of what principles govern the geometry of an animal’s internal representation as its knowledge set of latent patterns grows.

### Organization of clusters within individual contexts

We first evaluated whether, within an individual context, the arrangement of rACC neural activity clusters reflects how an animal executes a given sequence. Since successful execution of a sequence matching the target pattern requires producing ‘L’/’R’ symbols in the correct order, we investigated whether neural clusters would be organized to reflect this progression (*46, 57*). Indeed, such an ordered arrangement was apparent even in the space of the first three PCs (Fig. 4A). To quantify this, we searched for a direction in population activity space — i.e., weighted sum of activity of different neurons — that best tracked sequence progression. Focusing first on the minimal pattern matches, we used proportional odds logistic regression to find this ordinal direction. We then mapped the projected activity onto the corresponding sequence step and tested classification performance on held-out data (Fig. 4A inset; see Materials and Methods). Decoding of sequence progression from rACC activity was significantly above chance, even when the analysis was restricted to the first three PCs (Fig. 4B; average decoding accuracy of 0.97, 0.85, 0.62, and 0.58 for sequences of length 3, 4, 5, and 7, respectively, vs. chance levels of 0.33, 0.35, 0.2, and 0.14, p< 10^-5^, t-test). Qualitatively similar results were obtained when activity was aligned to side port (rather than center port) entries (average decoding accuracy: 0.96, 0.84, 0.63, 0.57 p< 10^-6^ vs chance, t-test). Progression decoding was even more accurate when the readout direction was computed in the high-dimensional activity space (Fig. 4B, average decoding accuracy: 0.98, 0.9, 0.75, 0.68 p< 10^-7^ vs chance, t-test), confirming that the ordering is not an artifact of dimensionality reduction. Moreover, decoding accuracy dropped substantially when cluster labels were randomly shuffled (fig. S4), indicating that the ordinal structure we observed is not just a byproduct of easier separability in high-dimensional spaces (*58*). These findings suggest that rACC does not just encode discrete states but organizes them in a neural trajectory that reflects the temporal order of steps in the learned pattern.

**Figure 4.**
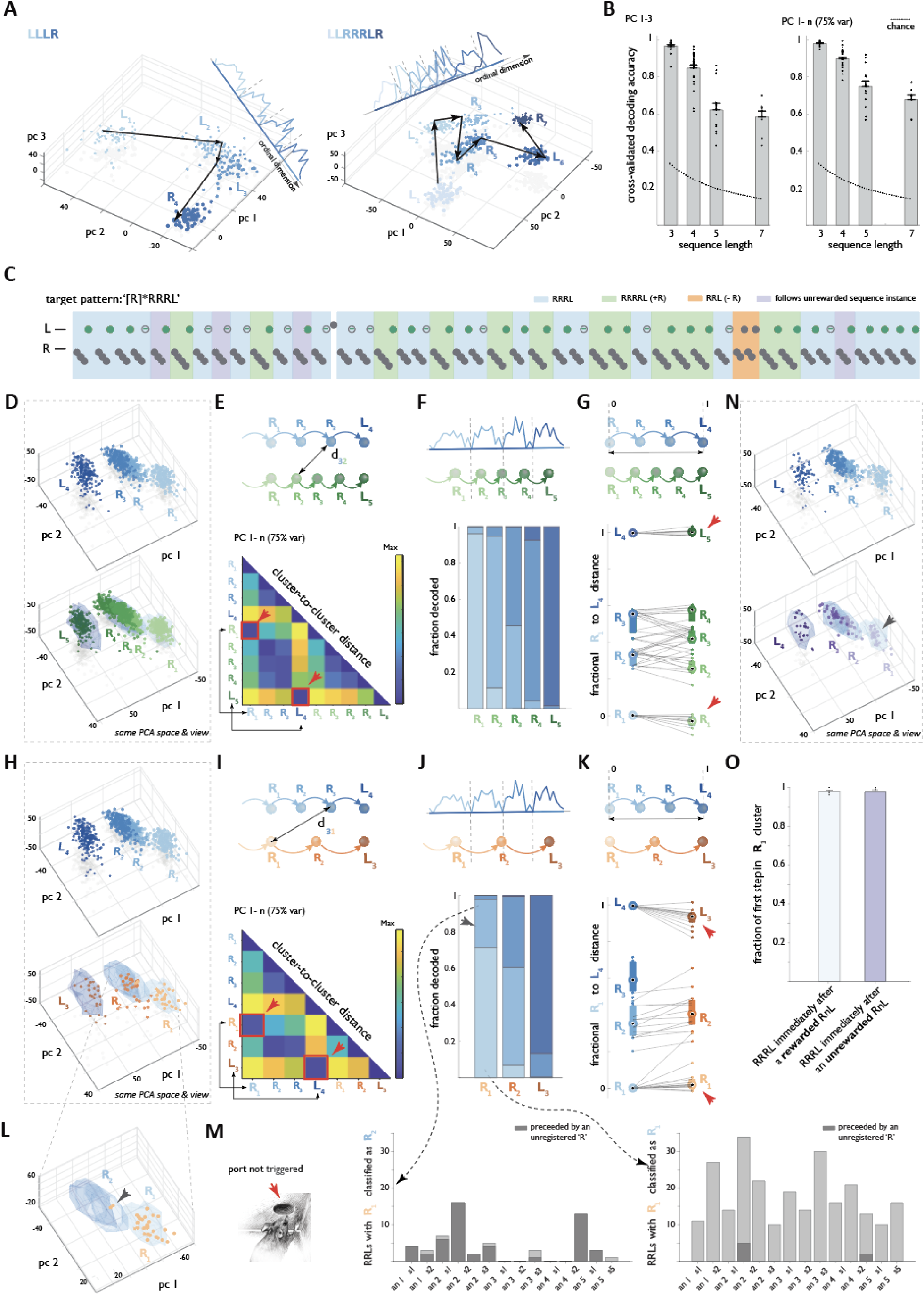
Within each context, rACC represents a bounded neural graph comprising a self-organized, editable set of sequentially organized activity nodes. (**A**) Example neural geometry from two sessions with distinct latent target patterns highlighting the representation of ordinal progression. Activity clusters are colored by sequence step, with color progressively saturating deeper into the sequence. Arrows connect activity patterns for steps in one sequence instance. (**B**) Performance of a linear ordinal position classifier on held-out data in the space of the first three principal components (left panel) and in high-dimensional space (right panel) for sessions with target patterns of different lengths. Dotted lines: chance performance. [n= 18,25,10,9 latent contexts, n= 9,14,10,9 sessions, N=3,4,3,4 animals for latent structures of order 3,4,5 and 7]. (**C**) Example behavioral trace for ‘RRRL’ context highlighting minimal pattern matches (blue), expanded matches (green) and pattern contractions (orange). Also highlighted are pattern matches that follow a single unrewarded sequence instance (purple). (**D**) Neural geometry for minimal pattern matches (‘RRRL’s, top panel) and for expanded pattern matches (‘RRRRL’s, plotted on the background of ‘RRRL’ cluster outlines in blue, bottom panel) for an example session. (**E**) Top panel: schematic of distances between cluster centroids. Bottom panel: d_ij matrix of cluster-to-cluster distances for the example in (D). Note that although the ordinal progression in the example PCA space is from right to left, the schematics and quantification here and in the rest of the figure use left-to-right convention. (**F**) Top panel: Schematic of applying a linear ordinal position classifier trained on the four steps of the ‘RRRL’ dataset to the five steps in ‘RRRRL’s. Bottom panel: Flattened confusion matrix for the schematized classification. Note the near-perfect classification of the initiating and terminating nodes. [n= 9 sessions; N=3 animals]. (**G**) Top panel: schematic for establishing fractional locations of the five ‘RRRRL’ graph nodes along the arc between the initiating and terminal nodes of the ‘RRRL’ graph. Bottom panel: Fractional location of the five ‘RRRRL’ activity nodes. Note that the first and the last clusters do not change location despite a different number of accommodated steps (red arrowheads). [n= 9 sessions; N=3 animals]. (**H,I**) same as in (D,E) but for ‘RRL’ target pattern contractions. (**J**) Same as in (F), but for ‘RRL’ pattern contractions. Note partial misclassification for step 1 in ‘RRL’ (arrowhead). [n= 14 sessions; N=5 animals]. (**K**) Same as in (G), but for ‘RRL’ pattern contractions. (**L**) A zoomed in portion of the PCA space from (**H**) focusing on R1 and R2 cluster locations for ‘RRRL’ (blue outlines). Arrowhead points to two R1 instances from ‘RRL’ dataset that map to second cluster from ‘RRRL’. Video analysis indicates these instances were preceded by an unregistered R step. (**M**) Distribution of all putative ‘RRL’ pattern contractions preceded by an unregistered ‘R’ step (left panel, schematizing port entry too shallow to trigger detection, see also fig. S5) displayed separately for ‘RRL’ instances with R1 mapping to cluster 2 of ‘RRRL’ (middle panel) and for ‘RRL’ instances with R1 mapping to cluster 1 of RRRL (right panel). See also fig. S5 for correction to confusion matrix and fractional cluster position following correction for unregistered sequence steps. (**N**) Neural geometry, in the same example session, of minimal target pattern matches (‘RRRL’s) that follow either a rewarded (top panel) or an unrewarded (bottom panel) sequence instance. (**O**) Fraction of behavioral steps immediately following rewarded and unrewarded sequence instances that map to initiating node of ‘RRRL’. [n= 9 sessions; N=3 animals].

We next examined how the geometry of rACC neural activity clusters is altered when animals edit their cognitive graphs, potentially to evaluate whether a missing state should be added or a redundant state eliminated (Fig. 1C,D). Focusing on animals performing in the ‘[R]*RRRL’ latent pattern context, we compared the geometry of the minimal (‘RRRL’) and expanded (‘RRRRL’) pattern matches (Fig. 4C). Despite differences in produced sequence length, cluster locations remained remarkably consistent for structurally aligned pattern expansions (Fig. 4D, E). Strikingly, the first and the last nodes typically maintained stable positions in the graph regardless of sequence length (Fig. 4G, location of first cluster −0.04 +/−0.01 vs 0, last cluster of 1.00+/−0.01 vs 1 for ‘RRRRL’ vs ‘RRRL’ sequence variants), whereas the number and position of intervening nodes was adjusted to accommodate the differing number of steps in the evaluated sequence variant (Fig. 4G). A similar pattern was observed for pattern contractions (‘RRL’s, Fig. 4H-K, location of first cluster 0.04 +/−0.02 vs 0, last cluster of 0.93+/−0.01 vs 1 for ‘RRL’ vs ‘RRRL’ sequence variants), though the first cluster stability appeared reduced (arrowhead in Fig. 4K). These results suggest that sequence variants could reflect intentional explorations of the state space, with the first and last nodes anchoring a stable frame of reference, while intermediate states are dynamically adjusted.

We also leveraged the crystalline structure of rACC activity to identify a subset of ‘RRL’ sequences, in which the first ‘R’ did not map to the expected anchoring cluster but to the second cluster of ‘RRRL’ instead (Fig. 4 J,L). We wondered if this mismatch reflected a discrepancy between the animal’s internal assessment of its sequence production and our experimental record of its progression. Possible explanations for such a mismatch might include lapses in memory, sequence execution errors, or shallow/quick port entries that went unregistered by our system. Indeed, when we examined video recordings for attempted port entries, we found that an unregistered right-port entry (Fig. 4M, left, see also fig. S5A) preceded over 89% of misaligned ‘RRL’ instances (Fig. 4M, middle), compared to only 3% in correctly aligned ‘RRL’ ones (Fig. 4M, right). Further, when ordinal step labels in the misaligned ‘RRL’ instances were reassigned to the animal’s likely internal assessment of ordinal position based on video analysis, the stability of the anchoring first cluster position improved significantly (fig. S5). Thus, neural activity states followed the animal’s internal model, even when our behavioral tracking system failed to capture it. As such, what appeared to be shortened misaligned ‘RRL’s were actually mis-registered ‘RRRL’s, while ‘RRL’s that started in the anchoring, R1 node of ‘RRRL’ graph represented true exploratory target pattern contractions. Notably, such deliberate target pattern contractions by expert animals would not be associated with the same reward expectation, suggesting that reaching the graph’s terminal node is a self-imposed goal. Combined, these analyses highlight the power of instantaneous state inference enabled by the crystalline nature of the rACC representation and show how differences in neural geometry across variations of a sequence can reveal animals’ internal punctuation of action sequences, including initial and terminal sequence steps.

Finally, we asked whether animals could re-start navigating their internal state spaces without using external reward for guidance. The pattern contractions, which were not rewarded, as well as target matches for which reward was intentionally withheld (Fig. 4C), offered an opportunity to study neural activity evolution past the terminal neural graph node in the absence of such guidance. Consistent with earlier behavioral findings (Fig. 1E) and substantiating that reaching the terminal node was a self-imposed goal, neural activity for behavioral symbols immediately following unrewarded sequence instances reliably returned to the first anchoring node (in 99 ± 1% of cases; Fig. 4N,O). Thus, reward is not required for the animal to restart state traversal within its internal model of the sequence. Since transitions between internal states are also necessarily independent of sensory feedback in this paradigm, these findings argue that within-context rACC representations constitute self-organized, ordered, and editable cognitive graphs of unguided latent structure discovery. How then are the individual neural graphs positioned with respect to one another as animals encounter contexts with different latent patterns?

### Across-context cluster geometry

Recent studies have suggested that the brain may support knowledge organization by mapping distances in conceptual or semantic space onto distances in neural activity space (*59, 60*). To test whether this principle applies to rACC representations of cognitive sequences, we examined whether distinct neural graphs — each corresponding to a different latent sequence structure — are organized within the overall geometry of the cluster space in a way that reflects their putative semantic similarity to the animal (Fig. 5A). We focused on three latent target patterns: ‘LLLR’, ‘RRRL’, and ‘RRRLR’, chosen to test distinct hypotheses about the type of across-context similarity that might be encoded. One possibility is that the organization reflects more proximal ‘explanatory power’ for reinforcement received when a latent pattern is first encountered, for example, based on the side at which reward is delivered. Under this scheme, neural graphs for ‘LLLR’ and ‘RRRLR’ would be grouped together (both predicting reward on the right side), and distant from ‘RRRL’, which corresponds to left-side reward delivery. Alternatively, the grouping might reflect a more general computational principle. One such principle is compositionality — the idea that complex knowledge structures can be built by flexibly combining simpler components, with the meaning of the whole being determined by its parts (*61*). In this framework, neural graphs for ‘RRRL’ and ‘RRRLR’, which share overlapping substructures, would be expected to occupy neighboring positions in cluster space. A third possibility is that across-context organization reflects higher-order abstraction, such as algebraic structure (*43*). For example, both ‘LLLR’ and ‘RRRL’ share an ‘AAAB’ pattern, and might therefore be grouped together, despite differing in constituent symbols. By examining the geometry of the cluster space across these three contexts, we sought to determine whether neural representations reflect any of these similarity metrics — reward location-based, compositional, or higher-order abstraction.

**Figure 5.**
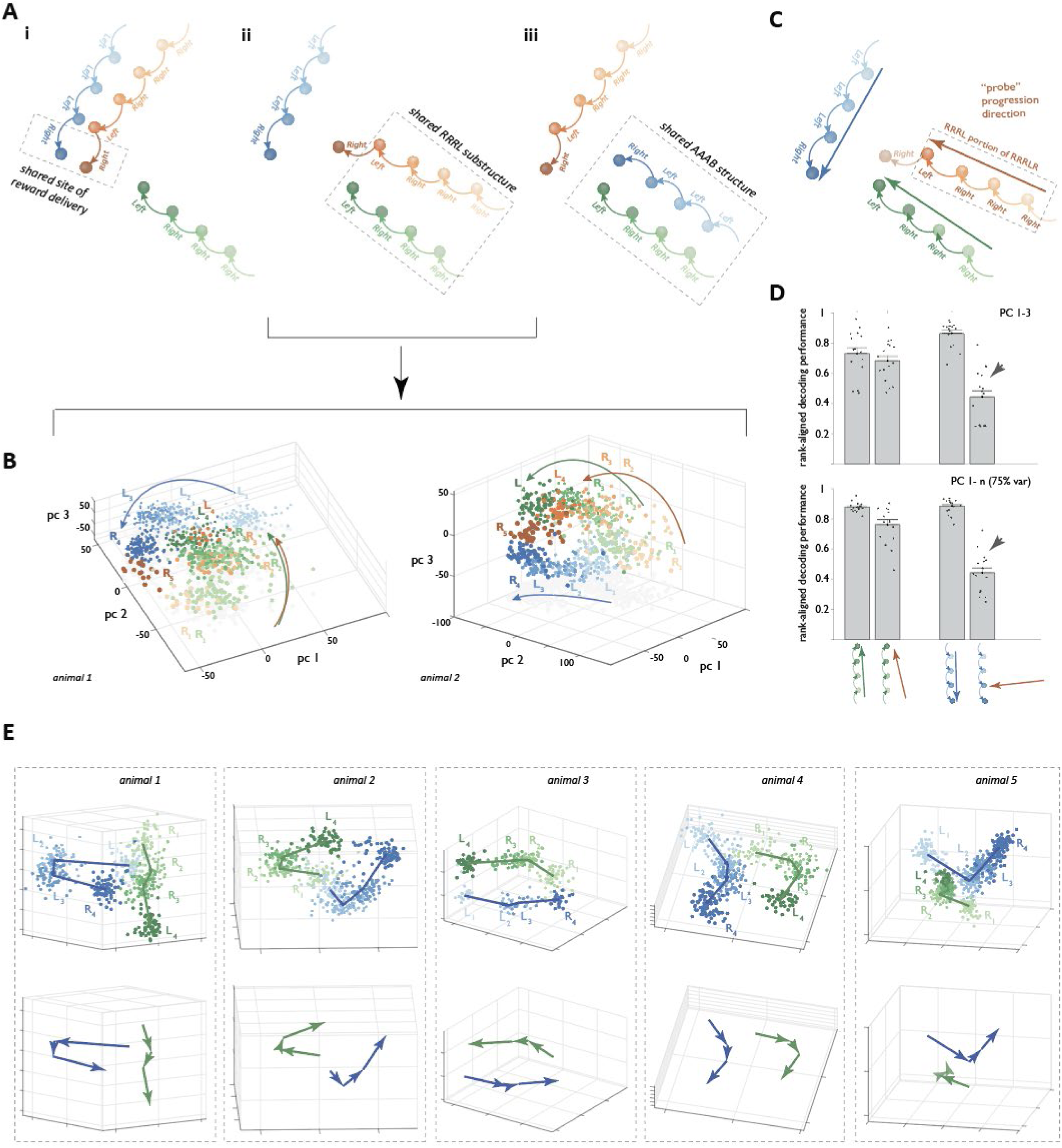
Alignment of neural graphs for sequences with shared sub-structures. (**A**) Schematics of possible groupings by shared site of reward delivery (‘R’ in ‘LLR’ and ‘RRRLR’) (i), shared sub-structure (‘RRRL’ in ‘RRRL’ and ‘RRRRLR’) (ii) and shared algebraic pattern (‘AAAB’ in ‘RRRL’ and ‘LLLR’). (**B**) Example neural geometry from two sessions plotted in the space of the first three principal components; activity clusters in each sequence are colored by the sequence step, with color progressively saturating deeper into the sequence. Note that a shorter analysis window was used (left panel) to separate the effect of terminal cluster movement upon center port exit (see Fig. 6 below). The arrow schematics note an alignment of green and orange graph arcs. (**C**) Schematic of the approach used to quantify graph alignment. (**D**) Left bars: Rank-aligned ordinal position decoding for ‘RRRL’ clusters (green nodes) using either their native ordinal dimension (green arrow in C and D), or one determined by the ‘RRRL’ component of the ‘RRRLR’ graph (orange arrow in C and D); Note the good decoding for ‘RRRL’ using the ordinal dimension for the corresponding substructure within RRRLR, both for analyses carried out in the space of the first three principal components (top panel) or in high dimensional space (bottom panel). Right bars: Ordinal position decoding for ‘LLLR’ clusters (blue nodes) using either their native ordinal dimension (blue arrow) or one determined by the ‘RRRL’ component of the ‘RRRLR’ graph (orange arrow); Note the markedly worse performance in decoding ‘LLLR’ using the ordinal dimension for the algebraically similar ‘RRRL’ sequence (arrowhead).. [n=15 sessions, N=5 animals]. (**E**). Alignment of neural graphs for ‘LLLR’ and ‘RRRL’ across 5 animals. Top panel: activity patterns for individual sequence instances are shown, along with the position of cluster centroids (thick lines). Bottom panel: cluster centroids only. Note the variation in alignments between clusters for these algebraically similar sequences.

In all five animals tested on within-session unsignalled changes between ‘LLLR’, ‘RRRL’, and ‘RRRLR’ latent contexts, we observed that the ‘RRRL’ and ‘RRRLR’ graphs progressed along aligned arcs, while the ‘LLLR’ graph followed a distinct trajectory, unique to each animal (Fig. 5). To quantify the geometric relationships among these representations, we used the ‘RRRLR’ sequence to define a sequence-progression direction for its embedded ‘RRRL’ substructure (Fig. 5C). We then assessed how well this direction could generalize to capture sequence progression in the ‘RRRL’ and ‘LLLR’ sequences. The underlying intuition is that if two sequences share an aligned structure in neural space, then the progression direction extracted from one should accurately decode progression through the steps of the other. Conversely, misaligned sequence representations would each yield poor decoding performance when using a direction derived from the other. We note that decoding performance is insensitive to vector sign—that is, the decoder is blind to whether it tracks forward or backward progression. To address this, we imposed a rank-direction constraint, ensuring that decoding reflected natural forward sequence order (see Materials and Methods). Further, for consistency, we included only trials where animals provided strong behavioral evidence of correctly identifying the latent pattern (see Materials and Methods). Using this approach, we found that the progression direction derived from ‘RRRLR’ generalized well to ‘RRRL’, but poorly to ‘LLLR’ (Fig. 5D). Specifically, decoding accuracy for ‘RRRL’ using the ‘probe’ direction (from ‘RRRLR’) was 70.5+/−3.5% in the space of first three principal components (76.2+/−3.2% in high dimensional space), compared to 72.1+/−3.5% (87.8+/−0.8%) using its native direction; for ‘LLLR’, the corresponding errors were 39.9+/−4.6% (44.3+/−3.3%) and 83.4+/−2.2% (88.5+/−1.3%), respectively (p< 10^-3^, Wilcoxon ranksum test, n=15 sessions, N=5 animals). Similar results were observed when analysis was anchored on side port entry rather than center port, further supporting the robustness of this effect (side-port decoding errors: 57.5+/−2.8% (68.7+/−3.3) vs. 62.2+/−3.2% (86.1+/−0.6%) for ‘RRRL’, 36.2+/−3.4% (37.7+/−2.9%) vs. 83.3+/−1.5% (88.9+/−0.9) for ‘LLLR’, p< 10^-3^, Wilcoxon ranksum test, n=15 sessions, N=5 animals). This alignment between ‘RRRL’ segments in the ‘RRRL’ and ‘RRRLR’ state spaces is unlikely to result from shared motor trajectories, as shown by the differing intermediate state geometry *within* these two graphs (see below). Instead, these results suggest that the geometry of the rACC state space reflects semantic similarity between ‘RRRL’ and ‘RRRLR’. By contrast, the distinctive embedding of ‘LLLR’ that differs across animals (Fig. 5E) indicates that unrelated sequences are not anchored to common subspaces, arguing that when a relationship to previously learned latent structures cannot be discerned, the new graph can be positioned flexibly. Given that a new graph should also progress towards an anchored goal state (Fig. 4), the observed flexibility raises the question of whether any additional organizational principle governs the placement of each graph’s anchoring nodes.

### Context-independent ‘sequence termination’ goal state

In guided experimental settings, goal states are often explicitly reflected in differential patterns of neural activity for cues signaling impending reward (*44, 62–65*). This led us to ask what kind of internal goal representation, if any, exists in our unguided setting, where animals discover latent structures without external instruction. Although the reward in each context is explicitly mapped to a specific side port (left or right), this mapping is not unique, since each port may be visited multiple times during a single sequence. Thus, physical location alone cannot specify the goal. Instead, based on our earlier findings (Fig. 4), the final node in each sequence graph may serve as an internal, context-specific representation of a goal: these terminal nodes displayed anchoring properties, and transitions out of them were reward-independent. We then asked whether a more abstract, context-independent goal node might be present—one that can help orient the cognitive graph of a new latent structure unrelated to previously encountered ones (Fig. 5D). To test this, we turned to the dataset with ‘LLLR’, ‘RRRL’ and ‘RRRLR’ latent contexts, where two alternative predictions can be distinguished (Fig. 6A): (i) If reward side defines the goal, terminal nodes should align according to left vs. right port outcomes. (ii) If an internal context-independent goal exists, terminal nodes should converge across sequences, despite different physical actions of producing the terminal symbols. We first examined the terminal clusters corresponding to rewarded executions of ‘LLLR’, ‘RRRL’, and ‘RRRLR’ (same subset used in the cross-projection analyses, Fig. 5). We found a marked convergence of these graph nodes, particularly when neural activity was aligned to side port entry (Fig. 6B). This convergence could reflect a common sensory-motor experience tied to reward delivery and consumption, or it could reflect an internal computational strategy in which the terminal state representation of each context converges onto a goal node shared across contexts. To distinguish between these possibilities, we evaluated the effect of broadening our analysis to include all sequences aligned with these target patterns, both rewarded and unrewarded (Fig 6Ci,ii) or limiting to just the unrewarded sequence variants (Fig. 6Di,ii). We found similar terminal node convergence across these datasets, arguing against the idea that convergence merely reflects external reward-driven input. Instead, this indicates an internally driven convergence towards a context-independent goal state across sequence terminations. Notably, this convergence appears to be facilitated by a reorganization of intermediate node geometry. Specifically, in the ‘RRR**L**R’ sequence, the non-terminal ‘L’ diverged away from its cognate but terminal ‘L’ in the ‘RRR**L**’ sequence (Fig. 6E, lighter orange L4 cluster diverging away from the green L4 cluster). Concomitantly, the terminal ‘L’ in ‘RRR**L**’ shifted toward the two terminal ‘R’ clusters (from ‘LLL**R**’ and ‘RRRL**R**’) (Fig. 6E, green L4 cluster shifting towards blue R4 and dark orange R5 clusters), despite substantial differences in ‘L’ and ‘R’ movement kinematics and spatial position between left and right side ports.

**Figure 6.**
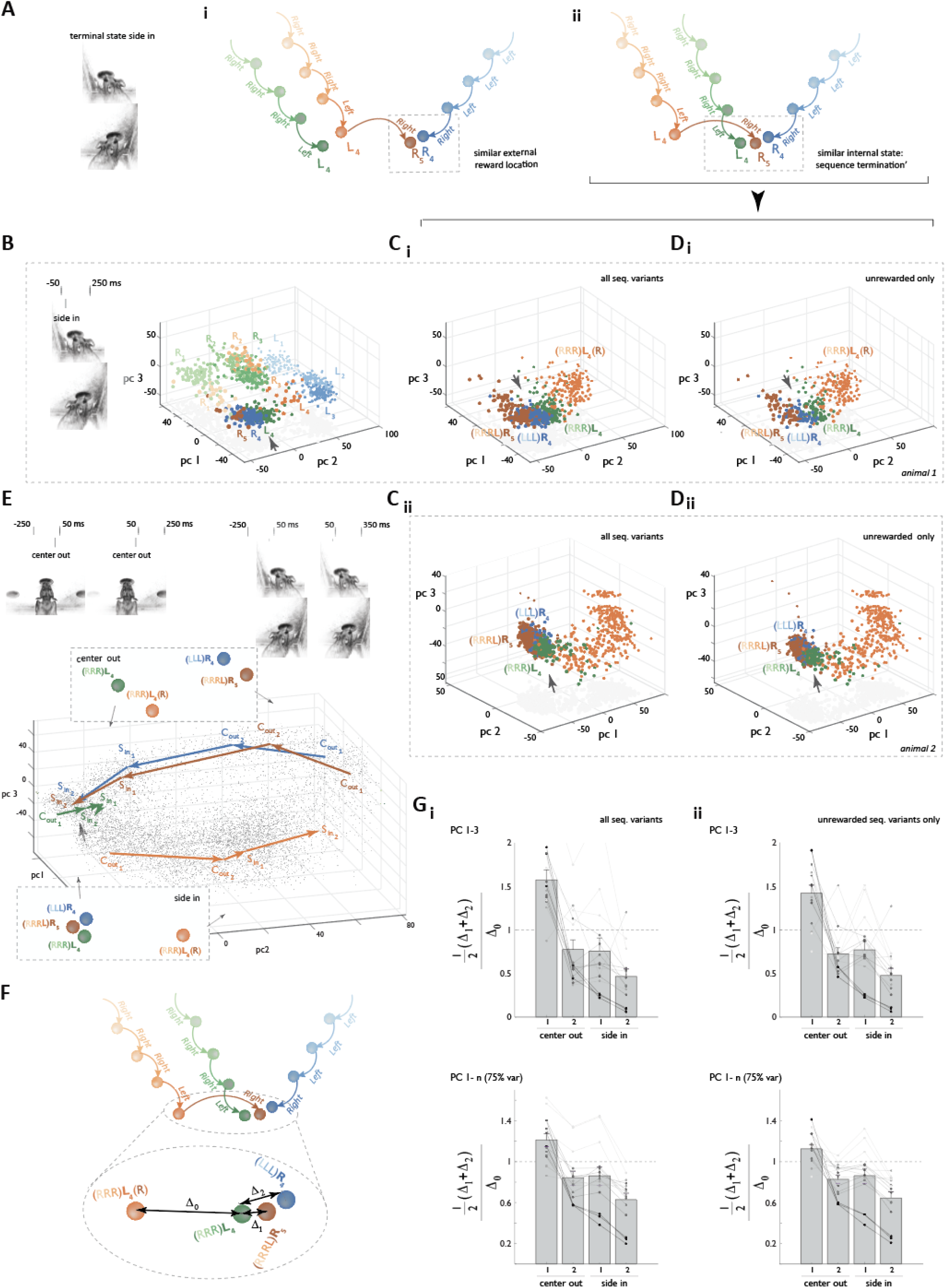
Context-specific neural graphs converge on a context-general goal node. (**A**) Schematics of two possible goal state configurations across contexts. Context-specific terminal states could be grouped by external reward location at left versus right port (i), or oriented to create a context-independent internally specified terminal state (ii). (**B**) Neural geometry for one of the example sessions from Figure 5, plotted in the space of the first three principal components of population activity at the time of entry into the side choice port (left panel); activity clusters are plotted for the sequences RRRL (green), LLLR (blue), and RRRLR (orange), with color progressively saturating deeper into each sequence. Only those sequence instances are shown that were included in Fig. 5 (see Materials and Methods). Note convergence of three terminal clusters (arrowhead) corresponding with schematic model Aii. (**C**) Neural geometry of the 3 terminal clusters and the cluster corresponding to the non-terminal ‘L’ in ‘RRR**L**R’ (that is motorically cognate to the terminal ‘L’ in ‘RRR**L**’). Data are plotted for just these 4 clusters in panel Ci for the same rat as in B, and for an additional rat in panel Cii. All sequence variants are included. (**E**) Neural geometry of cluster centroids from (C), plotted in the space of the first three principal components of an expanded activity space derived from 4 analysis windows (two windows anchored on center port exit and two on side port entry, top panel). Note the convergence of three terminal clusters between center port exit and side port entry (arrow). Boxed schematics illustrate the relationship between cluster centroids during at these center-out and side-in time points. (**F**) Quantification schematic. Distances from the terminal ‘L’ in ‘RRRL’ (green) to the other two terminal clusters (Δ_1_ and Δ_2_) are compared to the distance from the same ‘L’ and its motorically cognate but non-terminal ‘L’ in ‘RRRLR’ (Δ_0_). (**G**) Terminal state convergence metric (½( Δ_1_+ Δ_2_)/ Δ_0_) across the four analysis windows schematized in (E) for all sequence variants (Gi) or only unrewarded sequence variants (Gii) and calculated either in the space of first three principal components (top row) or in the high dimensional activity space (bottom row). Line shading differs by animal. [n=15 sessions, N=5 animals].

To determine more precisely when the neural representation shifted to bring terminal states into alignment, we examined trajectories associated with these four clusters at a finer temporal scale, by subdividing each ‘L’ and ‘R’ step into four consecutive analysis windows (Cout1, Cout2, Sin1, Sin2), beginning 250ms before withdrawing from the center port and ending 350 ms after entry to the side port (Fig 6E). By expanding the representation of each step into these within-step trajectories it was apparent that the realignment of the terminal ‘L’ in ‘RRR**L**’ with the terminal ‘R’ clusters in ‘RRRL**R**’ and its divergence from the cognate but non-terminal ‘L’ cluster in ‘RRR**L**R’ began as the animals exited the center port on the final step (Cout2) and culminated in convergence as they completed entry into the L or R side port associated with the step (Sout2). To quantify this pattern, we used a metric designed to capture relative cluster proximity (Fig. 6F), which documented a similar pattern of rapid, within-step convergence across multiple sessions and animals (Fig. 6G, fractional distance across all variants of 0.47+/−0.08 vs. 1.57+/−0.11 in the space of the first three principal components (0.63+/−0.06 vs 1.21+/−0.06 in high-dimensional space) for cluster arrangement after side port entry (Sout2) vs before center port exit (Cout1), p < 10^-5^, Wilcoxon ranksum test, n=15 sessions, N=5 animals). Thus, neural graphs representing different latent patterns converge at a common goal node, suggesting a computational strategy that organizes the structure of animals’ internal models as their knowledge grows with experience.

### Leveraging crystalline rACC representation for algorithmic insight

The experiments above suggest that rACC neural activity can be used for real-time readout of an animal’s interpretation of the environment (Fig. 4M). This could be especially powerful in situations where existing algorithmic models of behavior fall short in explaining individual behavioral patterns. To illustrate this point, we reexamined neural activity structuring in the ‘LLR’-‘RRL’ forced pattern alternation regime, which we used previously to scale latent structure complexity (Fig. 2). In this context, both ‘LLR and ‘RRL’ are reward-eligible, but must alternate for continued reinforcement (see above and Materials and Methods). This is in contrast with current models of behavior that typically assume that a rewarded behavioral pattern will remain so for some time (*66*). Nonetheless, all animals successfully discovered the alternation rule and maintained a high reward rate by flexibly switching between the two patterns. Since no constraints were placed on what behaviors could be interposed between ‘RRL’ and ‘LLR’ pattern matches, the resulting behavior was noticeably more heterogeneous within sessions, between sessions, and across animals than in contexts with simpler latent structure (data not shown). However, within this heterogeneity, one recurring sequence motif stood out across animals ‘RRL+RRLLR’ and its mirrored variant, ‘LLR+LLRRL’, with ‘+’ indicating reward delivery (Fig. 7A). The post-reward fragment of the motif, e.g. ‘RRLLR’, is particularly intriguing. As this experiment was done without feedback delays that enforce sequence punctuation, this fragment, could be interpreted in multiple ways, e.g., RRLxx, xRLLx, or xxLLR, each with its own algorithmic interpretation of how animals plan and organize their actions. Such ambiguity lies at the heart of the challenge in constructing algorithmic models of behavior in complex, unguided settings. An independent window into the animal’s parsing of its behavior would therefore be invaluable for such models. Indeed, despite the heterogeneity of animals’ behavior in this setting, rACC activity remained highly structured (Fig. 2C), with six stable clusters corresponding to the individual steps of the rewarded ‘RRL’ and ‘LLR’ sequences (Fig. 7B). We asked whether these clusters could help resolve the algorithmic ambiguity inherent in the observed intervening symbol motifs.

**Figure 7.**
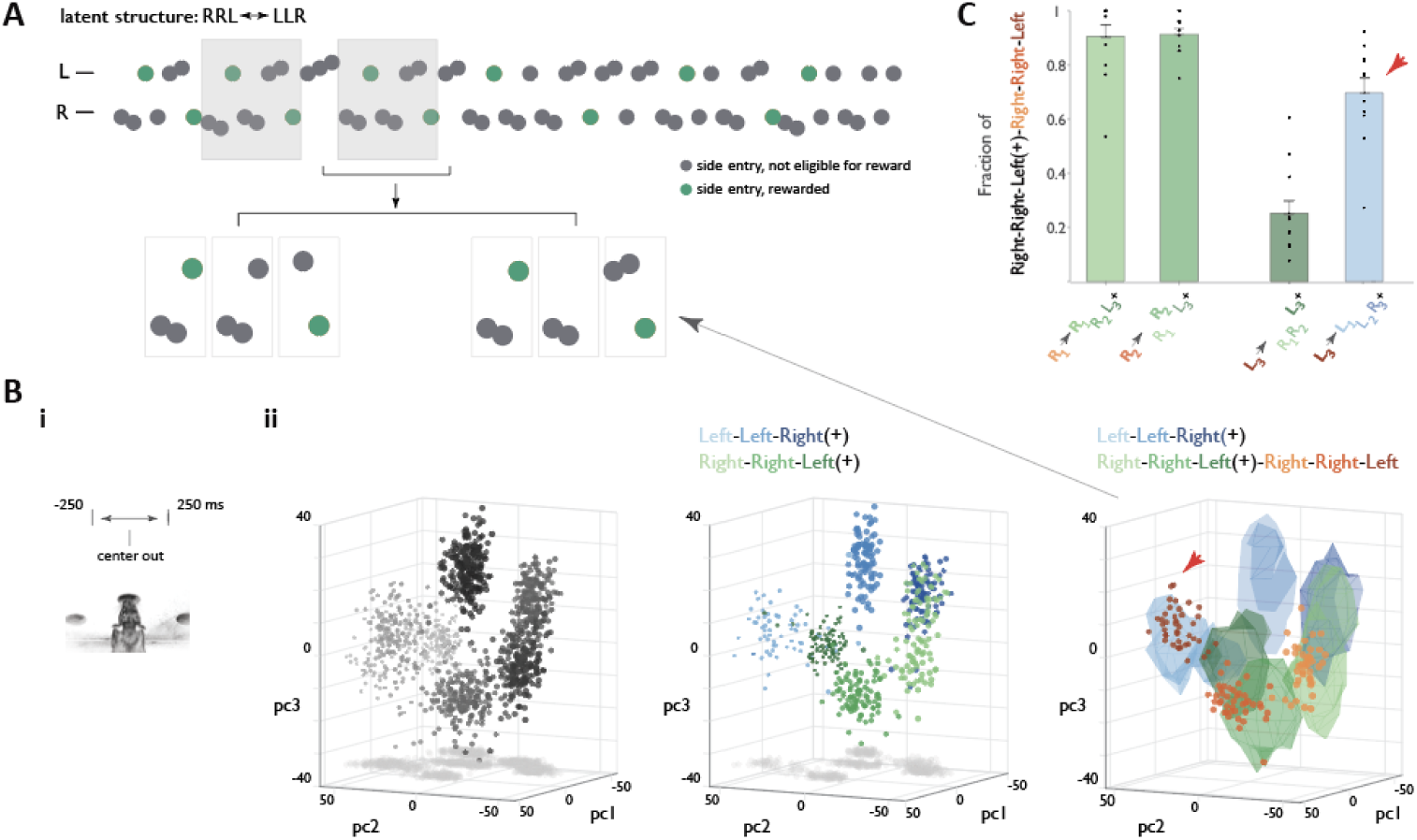
Leveraging rACC representation for algorithmic interpretation. (**A**) Example behavioral trace from a latent context requiring alternation between ‘LLR’ and ‘RRL’ patterns. Grey boxes highlight ‘RRL(+)RRLLR’ motif. (+): reward delivery. (**B**) Neural geometry at the time of center port exit (i) in forced pattern alternation context for an example session, plotted in the space of the first three principal components for the entire session. Left panel: all behavioral steps in this context (shading used here solely to facilitate visualization of depth). Middle panel: ‘LLR’(+) and ‘RRL’(+) instances, colored by individual step. Right panel: unrewarded ‘RRL’ instances that immediately follow rewarded ‘RRL’s, colored by individual step and overlayed on the contours of the 6 clusters from the middle panel. Note that the majority of points corresponding to the terminal L in ‘RRL(+)RR**L’** (dark orange, arrowhead) fall on a blue contour, corresponding to L1 in ‘**L**LR’(+). (**C**) Fraction of points (activity patterns in individual motif instances) for each of the three steps in ‘RRL(+)**RRL’** aligning with the cognate clusters in ‘RRL(+)’ as well as fraction of ‘RRL(+)RR**L’** points aligning with L1 in ‘**L**LR’(+) (red arrow).

We compared rACC activity at center port exits — before the animal knew if it would be rewarded — for individual steps of the first and second ‘RRL’ in the ‘RRL+RRL’ motif. The first two steps (the two ‘R’s) of the second ‘RRL’ consistently mapped to the clusters expected from the mapping established by the first ‘RRL’ (in 11 out of 11 sessions from 3 animals; overall number of R1 and R2-mapping ‘RRL+**RR**L’ *instances* across sessions: 91+/−4% and 91+/−2% respectively, Figs. 7B, 7C left two bars). Unexpectedly, the third step (the ‘L’) in the second ‘RRL’ did not map to the cluster associated with the ‘L’ of the first ‘RRL’. Instead, it reliably mapped to the cluster corresponding to ‘L1’ of rewarded ‘LLR’ instances (in 10 out of 11 sessions from 3 animals; overall number of L1-mapping ‘RRL+RR**L**’ *instances* across sessions: 70+/−5%, Figs. 7B, 7C right two bars). Thus, in contrast to a smoothing assumption of most behavioral models, these data suggest that the animal was treating the second ‘L’ symbol as the first ‘L’ of an ‘LLR’, so that the correct algorithmic parsing of the ‘RRL+RRLLR’ motif is ‘RRL’→‘RR’→‘LLR’. This result highlights the interpretational power of the prefrontal neural graphs that we have uncovered. Rather than just being a lower-dimensional representation of neural phenomena, they reflect cognitive graphs and thus can reveal an animal’s model of the world and current objective in ambiguous settings.

At the core of higher cognition is the ability to build behaviorally relevant structured knowledge from the complex stream of information that we gather from our interactions with the environment. Recent theoretical accounts have argued that frontal regions may be particularly important for creating a mental representation that focuses solely on states relevant to animals’ current objective (*8, 26*). The abstract graph-like nature of rat rACC population representations during unguided latent sequence discovery substantiates just such a role for frontal cortex. Its crystalline abstraction of animals’ experience and actions corresponds to the minimal states required to achieve objectives in the paradigm: compact enough for planning yet not so compact as to lose behaviorally relevant distinctions (*67*). In our paradigm of sequence production, edits to the animals’ neural graphs were in the number and position of activity clusters between terminal nodes that represent ‘start’ and ‘goal’ states. This stability of start and goal states within the rACC neural representation may allow animals to initiate active evaluation of distinct sequences while maintaining the significance of these critical anchoring states (*68, 69*). All sequence explorations would then have these defined starts and ends themselves, which would allow their neural graphs to be automatically integrated into the same schema if successful, before being actively refined to the minimal necessary set of nodes. Indeed, if such agency and active interaction with a dynamic environment is at the core of the prefrontal cortex’s role in the construction of cognitive graphs (*26, 67, 70–72*), our unguided behavioral paradigm may have been particularly well-suited to engaging this brain region and revealing its representational schema.

A key challenge in reinforcement learning and AI is the credit assignment problem— attributing reinforcement to a specific action or sequence of actions in the past (*73*). However, all solutions to this problem of cause and effect assume that the agent can rely on a representational schema with states that represent potentially relevant causes (*3*). In reality, generating appropriate states itself is a challenge that requires parsing rich, detailed, and continuous streams of information for relevant structure (*19, 74–76*). An additional complication when exploring new contexts is that this tentative parsing process must co-occur with credit assignment, implying that states are still evolving as they are being used to develop models of their significance for reinforcement. Indeed, in open-ended settings, arriving at a parsing that is compact enough to make reasoning and planning feasible yet detailed enough to capture behaviorally relevant distinctions is often the most challenging part of the problem (*4*). The unguided sequence discovery framework that we have deployed provides experimental access to this complex process, offering a window into efficient solutions that evolution has selected for these highly challenging, yet-unsolved computational problems.

## Acknowledgments

We are grateful to the incredible Janelia Vivarium team for pampering the furry stars of this story. We thank Mattias Karlsson and Spikegadgets for headstage customization; Andrea Gugiu, Jon Arnold, Bruce Bowers and Bill Biddle from Janelia Experimental Technology for help with hardware prototyping, fabrication and assembly; Lillian Tong for help with pilot delayed feedback experiments. We appreciate the many useful discussions with Josh Dudman, Marius Pachitariu, Ann Hermundstad, Sandro Romani, as well as members of the Koay and Tervo labs at various stages of this project. We are truly indebted to Vivek Jayaraman, Sue Ann Koay, Michael Brainard and Gerry Rubin for extensive discussions, advice and comments on the manuscript.

## Author contributions

Conceptualization: MM, DGRT, AYK

Methodology: MM, MP, HW, EK, AL, SD, DGRT, AYK

Investigation: MM, MP, HW, EK, RB, AYK

Analysis: MM, MP, HW

Writing – original draft: SD, DGRT, AYK

Writing – review & editing: MM, MP, HW, EK, AL, RB, SD, DGRT, AYK

## Data and materials availability

All data and code will be made available upon request.

## Materials and Methods

### Subjects

All experiments were done in male Long Evans rats (400-500g). Animals were kept at 85% of their initial body weight before food restriction and maintained on a 12hr light/12hr dark schedule. Experiments were conducted according to National Institutes of Health guidelines for animal research and were approved by the Institutional Animal Care and Use Committee at HHMI’s Janelia Farm Research Campus.

### Behavioral apparatus

All behavior was confined to a box with 23 cm high plastic walls and stainless-steel floors. The floor of the box was 25cm by 34 cm, and the custom-made nose ports (https://karpova-lab.github.io/nosepoke) were all arranged on one of the 25 cm walls. All hardware was controlled and monitored with a custom-programmed microcontroller running MicroPython (micropython.org), which in turn communicated via USB to a PC running a PyControl GUI ( https://github.com/pyControl/code). Nose port entries were detected with an infrared beam-break detector (IR LED and photodiode pair) or with a Time-of-Flight sensor. Successful port interaction was typically associated with one or more types of feedback: a change in the state of a port-embedded LED, a short auditory tone or port vibration. This was done to minimize the number of unregistered behavioral symbols arising from port interactions that were too shallow or too fast. Note, however, that our pilot ground truthing experiments established that omitting any feedback had little impact on the animals’ ability to discover latent structure.

Reward for eligible behavioral symbols was directly at the side ports. Reward for food-restricted animals was in the form of a small volume (typically, 0.1-0.25 ml) of 10% sucrose solution mixed with black cherry Kool-Aid. Reward was delivered with the help of a custom made syringe pump (https://karpova-lab.github.io/syringe-pump/latest/), and dispensed for eligible behavioral symbols by about 300 msec following side port entry, unless extra feedback delays were imposed (see below).

### Behavioral paradigm and training

Food-restricted animals learned to pair center and side port entries and discover latent sequential structure (an experimenter defined target pattern of ‘L’/’R’ symbols) with no explicit guidance. There was no step-by-step shaping, but early in training, the embedded latent structure was typically restricted to 3-4 symbol sequences. Animals were typically trained every other day, and were given about 4 hours in the behavioral box. Most sessions – including the very first ones – included unsignalled, block-wise changes in the structure of the latent target pattern. Block duration varied depending on the complexity of the latent target and the animals’ level of proficiency but typically ranged from 250 to 1000 behavioral steps. A particular ‘context’ comprised all blocks with a specific latent target pattern.

To encourage initial exploratory generation of symbol sequences, novel patterns appearing in the behavioral stream were rewarded in addition to target pattern matches during the first ∼500 successful center port-side port pairings, with the fraction of rewarded novel patterns decreasing gradually over that time frame. Throughout the rest of the animals’ experience in this paradigm, reward was delivered only when a match to the specific experimenter-defined latent target pattern appeared in the behavioral stream, and only after the last symbol was produced. If feedback delays were deployed, in sessions where animals were first challenged with feedback delays, feedback delay (with respect to side port entry) was slowly (in 1-2 msec increments) increased with every produced symbol until it reached 1-3 seconds late in the session. In behavioral probe sessions (Fig. 1H, fig. S1A) feedback delay was increased to the target level over the first 100 behavioral symbols. Feedback delay of 3 seconds was sufficient to elicit a robust pattern of preferential pausing at sequence ends in all animals. Note that feedback delays were not deployed during electrophysiological data collection to avoid the associated motor confound when evaluating the meaning of the terminal graph node.

When necessitated by the target pattern composition (e.g same start and end symbol, or pattern alternation setting), the sliding frame for matching behavioral symbol stream to target was explicitly reset upon encountering a pattern match.

A subset of sessions incorporated reward omission for 10-30% of target pattern matches. Over 90% of all animals became proficient with the first set of latent target patterns within 1-3 weeks (3-10 sessions) of training. After sufficient data was collected from animals proficient in the first set of patterns repeatedly discovering them in sessions with unsignalled block-wise changes in target pattern, animals were challenged to discover longer patterns, Longer target patterns were typically constructed either through pattern lengthening or pattern composition. To facilitate discovery of more complex patterns, as well as to assess the process of refining the internal model to the minimal number of necessary states, target patterns were often defined in the form of a regular expression, permitting flexible expansion of specific repeated units, indicated by [A]*, where A is a particular set of symbols.

Minimal matches to target patterns were between 3 and 7 symbols. The full set of target patterns – presented in their regular expression format – deployed at various stages of this study was: ‘[L]*LLR’, ‘[R]*RRL’, ‘[L]*LLLR’, ‘[R]*RRRL’,’[L]*LRLRR’, ‘[R]RLRLL’, ‘[LR]*LRLRR’, ‘[R]*RRRLR’, ‘[L]*LLRRL’, ‘[L]*LLR[R]*RRL’, ‘[R]*RRL[L]*LLR’, ‘[LR]*LRLRLRR’ and ‘[L]*LLR[R]*RRLR’.

### Behavioral analysis (Fig. 1D,E)

Prevalence of distinct sequence classes (Fig. 1D) was quantified by 1) categorizing each ‘L’ or ‘R’ symbol within individual sessions as belonging to an instance of a particular class, 2) computing a fraction of all symbols in the contextually-appropriate part of the session falling into each of the classes. ‘Minimal match to target pattern’ and ‘expanded match to target pattern’ were defined according to the specific regular expression form of the experimenter-defined latent target. For instance, for ‘[R]*RRRL’, ‘RRRL’ constituted the minimal match, while ‘RRRRL’ and ‘RRRRRL’ were possible expanded matches. In practice, permitted expansions of a repeated unit rarely exceeded two in expert animals. As such, the class of ‘expanded match to target’ for characterizing the structure of behavior in expert animals was limited to addition of 1 or 2 symbols. Pattern contraction was defined for target patterns 4 symbols and longer and were defined as a structurally aligned sequence missing one repeated symbol. Symbols were characterized as ‘matches to previous pattern’ if they were part of a sequential context embedding a match to a previously discovered pattern.

To characterize how robustly animals restart with the correct symbol of the target pattern with and without reward (Fig. 1E), we considered all the minimal and expanded target pattern matches, as well as pattern contractions. For each instance in this set, we calculated the gap between the last symbol in the current class instance and the first symbol in the next class instance. For each target pattern length and each session, we then calculated mean gap over all instances that did or did not result in reinforcement and summarized the data across sessions. Separately we calculated the probability of restarting with the correct first symbol of the target pattern as a probability of zero gap.

### Electrophysiological recordings

A total of 7 animals were implanted for collection of neural activity after the ninitial proficieny with latent sequence discovery was attained. Neuropixels 1.0 silicon probe (6 animals) or a microdrive array containing 16 independently movable tetrodes (1 animal) was chronically implanted on the head of the animal. Each tetrode was constructed by twisting and fusing together four insulated 13 μm wires (stablohm 800A, California Fine Wire). Each tetrode tip was gold-plated to reduce impedance to 200-300 kΩ at 1 kHz. Within the implant, the tetrodes converged to a vertically-aligned oval bundle (1 mm x 2 mmd). angled at 0° with respect to vertical (pointing towards midline after implantation).

For the probe / drive implantation surgery, trained animals were initially anaesthetized with 5% isoflurane gas (1.0 L/min) and mounted in a stereotaxic frame (Kopf Instruments). After 10-15 minutes, isoflurane was reduced to 1.5-2.0% and the flow rate to 0.7 L/min. A local anesthetic (Bupivacaine) was injected under the skin 10 minutes before making an incision. Small stainless steel bone screws and dental cement were used to secure the implant to the skull. Small stainless steel ground screw was placed above the cerebellum and connected to a wire leading to the system ground. A unilateral craniotomy (1.0 by 2.0 mm) was drilled in the skull above the site of recording and centered 3.5 mm anterior and 0.6 mm lateral to Bregma (right or left hemisphere) when recordings were targeted to the ACC.

For probe implant a single Neuropixels 1.0 probe was lowered to a depth of 6.0–6.3 mm from the brain surface. The probe / microdrive array was permanently fixed to the skull with C&B Metabond® Quick Adhesive Cement System (Parkell) and a protective enclosure attached to skull around the probe. For the Microdrive implant, before the animal woke up, all tetrodes were advanced into the brain ∼1.20 mm deep from the brain surface. Animals were allowed to recover from surgery for 10-14 days, over which time tetrodes in the one animal implanted with a Microdrive array were slowly lowered, moving approximately 40 μm/day on average.

After about 10 days, animals were re-acclimated to the behavioral paradigm. When motivation and proficiency of familiar latent pattern discovery regained pre-surgical levels, recording sessions began.

Data from all the animals were collected using the wireless headstage and datalogger (Horizontal Headstage 128ch with Datalogger, SpikeGadgets, https://spikegadgets.com/products/hh128-datalogger/ for tetrodes and Neuropixels Datalogger headstage, SpikeGadgets, https://spikegadgets.com/products/neuropixels-datalogger-headstage/ for the probe), powered by small Li-Polymer batteries affixed directly to the headstage. Battery rundown constrained the duration of recording sessions (typically, 3-5 hrs). Animals self-paced their behavior, and sometime took breaks (typically 5-30 minutes). Typically, electrophysiological data were collected for a stream of 1000-5000 behavioral symbols. For tetrode-implanted animal, recordings were made exclusively in rACC. For probe-implanted animals, 384 out of 960 channels were preselected to cover the depth of 0 to 4 mm from the brain surface to maximize the chance of picking up individual units in more ventral parts of mPFC. Channels close to the brain surface rarely had active units making it challenging to sample enough population activity in mPFC area dorsal to rACC.

### Ephys data preprocessing

Distinct data streams --behavioral event registration, time-of-flight signal from nose ports, video feed and electrophysiological data --were synchronized under a single clock. 4 key time stamps were established for each behavioral symbol: time of center port entry, time of center port exit, time of side port entry, and time of side port exit. If the IR beam was broken, or time-of-flight threshold exceeded, more than one time while the animal interacted with a particular port, the first timestamp in the series was taken for entries and the last time stamp for exits. No restrictions were imposed on the time between center and side port entries or on the detailed trajectory taken between the ports, thus some variation in the kinematics of symbol execution was present in all animals. Thus, with the exception of population trajectory visualization in Fig. 3 (see below), we analyzed population activity in defined fixed windows (500 msec in duration unless otherwise specified) aligned to port trigger events. This avoided the need for time warping when comparing neural activity between individual symbols. Because different animals displayed different center port dwell times, the center port-aligned analysis window was usually centered on port exit. In contrast, side port-aligned window was usually centered on port entry to exclude the period of reward delivery and consumption.

Once the analysis window was chosen, firing rates associated with all individual symbols in the continuous behavioral stream was calculated for all isolated units (by dividing spike number by the window length). The pre-processed data for each session was then assembled into a two-dimensional matrix with individual isolated units as rows and their FRs for individual behavioral symbols as columns.

### Ephys Data analysis

All analyses were performed using custom scripts written in MATLAB (MathWorks).

### Session selection

To ensure adequate sample size for electrophysiological analyses, we selected sessions with the highest prevalence of target pattern matches, aiming to include 3 sessions per animal for each type of latent context or context combination. When more than 3 sessions were available, we aimed to include sessions separated by days to weeks to confirm that the neural space geometry remained stable for each animal.

### Dimensionality reduction and population activity space

To visualize this high-dimensional activity while preserving its geometry, we applied principal component analysis (PCA), projecting the high dimensional population activity --defined by the matrix above --into the space of three dimensions that captured the most variance. Individual points on the resulting 3D plots thus represented population activity associated with a specific behavioral symbol. Population activity for all behavioral symbols produced over the course of a given session was used to construct the PC space for all analyses in the study. In contrast, the set of symbols for which population activity patterns were displayed in a given figure or used in a specific analysis varied depending on the question asked. Population activity plots and unsupervised analyses in Figs. 2 and 3 utilized the entire behavioral stream. In contrast, plots and supervised analyses of the geometry of target pattern matches (Figs. 3C-3, 4-7) focused on symbol constituents within matches to target patterns indicated.

### Clustering and fraction of variance left unexplained (Figs. 2, 3)

For unsupervised assessment of clustering, we used a standard iterative algorithm (k-means) that uses a distance metric to assign data points to one of k clusters, varying the values of k from 1 to 8. For each case, we calculated the amount of variance in the data left unexplained by a model with k clusters, as follows:

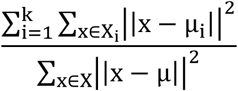

- *X* as the set of all data points
- *X*_*i*_ as the set of points in cluster i
- *μ* as the mean of all points
- *μ*_*i*_ as the mean of cluster i

These analyses were done, separately, either in the space of the first three principal components (PCs), or in the space of as many PCs --determined by examining the eigenvalues - as was necessary to capture 75% of total variance in the full firing rate matrix (7-20 PCs for different animals)

### Continuous trajectories (Fig. 3D, E)

To visualize continuous trajectories, raw spike time data for the entire session was binned in 10ms bins. The binned data was then smoothed by convolve it twice with an acausal half gaussian kernel (standard deviation of 150 msec), resulting in a complete neural trajectory for the session with a 10 msec resolution.

To visualize target pattern-specific trajectories in Fig. 3D, we defined – for each pattern match instance – ‘Side out’ of the preceding behavioral symbol as trajectory start, and ‘Side in’ on the last symbol of the pattern as trajectory end. This choice intentionally omits reward collection period. The binned trajectory data between these bracketing windows is then projected into the PCA space defined by the convention above in the data preprocessing section (first three principal components derived based on firing rates of all neurons in the 500 msec window centered on center port exit).

### Trajectory overlap (Figure 3E)

To estimate the extent to which trajectories associated with ‘LLR’ and ‘RRL’ pattern matches overlapped, we first constructed a PCA space that more broadly represented the population dynamics than the PCA spaces for a single analysis window centered on a specific port interaction. Instead, we effectively joined 4 high dimensional spaces: windows centered on Center port entry, Center port exit, Side port entry and Side port exit, and concatenated the four corresponding matrices of firing rates for individual neurons within each window. We then constructed a joint reduced space by performing PCA on the concatenated firing rate data, and taking the first three principal components.

We then binned this joint reduced PCA space into bins of 10 PC units. Unsing the smoothed trajectory datapoints above, we calculated the number of ‘LLR’ (‘RRL’) points in each PCA space bin. We then define ‘LLR’ (‘RRL’) trajectory subspace as the set of bins that covers 95% percentile of ‘LLR’ (‘RRL’) trajectory points. Having established per-context parcellation n of the PCA space, we then quantify the number of points from the other context that enters any bins in that space.

### Ordinal progression decoding (Fig. 4B)

To find the best direction in activity space for ordinal decoding without enforcing equal spacing between individual steps, we used proportional odds logistic regression. In separate analyses, we performed this search for the best direction by starting either with the space of the first three principal components, or with the space of as many PCs as was necessary to capture 75& of activity variance.

Once the best direction was found, we projected all training data points on that direction to obtain N distributions along that axis for neural activity patterns associated with an N-step sequence. We then determine ordinal classification boundaries using multi-class ROC classification:

1. Agnostic of ground truth ordinal information, we sort the training distributions according to their median location along the decoding axis
2. For each pair of adjacent distributions n and n+1, we find the boundary that maximizes the difference of the two cumulative distributions

Once classification boundaries were obtained from the training data, classification accuracy was tested on left-out test data.

Since many sessions had multiple context changes, we sought to eliminate periods of contextual uncertainty, restricting our analysis dataset to the set of minimal pattern matches that were preceded by reinforcement for successful target pattern matching. If the resulting sub selection left fewer than 10 sequence instances, the session was included from analyses.

For datasets with over 30 sequence instances, 80% of data was used for training and 20 % for testing. For datasets with more than 10, but fewer than 30 instances, -one-out cross-validation was done instead. In all cases, cross-validation procedure was repeated 10 times.

Chance level of classification accuracy was defined analytically as 1/N for an N-step sequence.

To verify that the ordinal structure we observed is not just a byproduct of easier separability in high-dimensional spaces we performed the following permutation control. For each session, we produced all possible re-indexing of the sequence steps (shuffled ordinal labels associated with individual clusters). For each reindexing, we find its best ordinal dimension and calculate decoding score without the more rigorous cross-validation step. These raw estimates are given in fig. S4A, with each permutation represented as a single point. We compare the performance of different permutations across sessions by expressing how well each permutation did in a particular session compared to others as a percentile and averaging those percentiles across all relevant sessions. Thus, for each target pattern length and each permutation, we acquire a single score: average percentile value that. We visualize the distribution of these values as violin plots (fig. S4B). The average percentile performance of the correct ordinal indexing variant (1,2,..N) is overlayed in green.

### Point cloud density estimation for cluster outline (Fig. 4D,H,N,L)

To visualize a cluster outline, kernel density estimation was performed using gaussian kernel with sigma 5 and then we visualize an isosurface corresponding to the 95th percentile.

### Analysis of exploratory graph editing (Fig. 4D-K)

Three animals displayed high frequency of exploratory graph editing, two in the ‘[R]*RRRL’ context, and one in ‘[L]*LLLR’ context. Analyses described below for the ‘[R}*RRRL’ case were mirrored wrt individual ‘R’/’L’ symbols for the third animal.

In each ‘[R]*RRRL’ session, analyses started with identifying a set of minimal target pattern matches, ie. ‘RRRL’s,. We then identified the associated ordinal progression dimension in the PC space that captured 75% of activity variance, and established classification boundaries for classifying the 4 ordinal steps of ‘RRRL’ as described above for Fig. 4B.

**<u>(4F)</u>** We next focused on the subset of expanded target pattern matches, specifically the set of all ‘RRRRL’s, and projected the associated activity patterns onto the ‘RRRL’ ordinal progression dimensioned. Using the ‘RRRL’ ordinal classification boundaries, we assigned each point from the ‘RRRRL’ dataset one of four classificiation labels (“R1”, “R2”, “R3”, or “L4”). Separating all ‘RRRRL’ points according to five groups THEIR true ordinal step. we quantified, within each group, the fraction of each of the four ‘RRRL’-based classification labels. The final plot in Fig. 4F displays average proportions across nine sessions from 3 animals.

**<u>(4G)</u>** For Figure 4G, instead of focusing on classification boundaries, we established the precise locations of the five clusters in ‘RRRRL’ dataset along the ‘RRRL’ progression dimension. For each session, the location of the “R1” cluster median in the ‘RRRL’ dataset was labeled as 0, and of the “L4” cluster as 1. Note that other clusters didn’t have to be between 0 and 1. Hence, we calculated the locations of “R2” and “R3” in ‘RRRL’ and of all five clusters in ‘RRRRL’

Analyses in Fig. 4F had established that “R1”, “R2”, “R4”, “L5” instances in ‘RRRRL’ were largely classified as “R1”, “R2”, “R3”, “L4” of ‘RRRL” respectively, whereas “R3” instances of ‘RRRRL’ were classified as either “R2” or “R3”, namely, the middle two steps of “RRRL”. The 6 lines in 4G (each connecting cluster medians within a single session) were chosen to reflect those relationships. Overlayed bars reflect the 25^th^, 75^th^, and 50^th^ percentiles across 9 sessions.

**<u>(4I-K)</u>** A conceptually similar approach to one in (4F,G) was used to analyze exploratory graph contractions from 14 ‘[R]*RRRL’ sessions across 4 animals.

### Analysis of unregistered port entries (Fig. 4M)

To determine if any behavioral symbols were missed due to port interactions that were too shallow or too fast, we carefully performed detailed trajectory reconstruction, using yolov8 video tracking model. Having extracted the location of a rat’s head frame by frame, we collected a set of labeled ‘at port’ examples and a set of labeled ‘out of port’ examples for classifier training. Classifier was trained classifier using an ensemble of boosted classification trees (fitcensemble MATLAB) and apply to detect nose port entry events that were visually similar to the registered ones but were not registered by the hardware. For analysis in Fig. 4M, we specifically checked if there were any unregistered ‘R’ symbols preceding each putative ‘RRL’ instance.

### Analysis of transitions out of the terminal node in the absence of reward (Fig. 4O)

For these analyses, we used the 14 ‘[R*]RRRL’ sessions. For analysis, we sub-selected minimal target pattern matches, i.e. ‘RRRL’s, that immediately followed a structurally-aligned sequence instances, i.e. an an instance of RnL (n>=2). We then divided that set into two groups, depending on whether the preceding RnL was rewarded or not. Using the classifier from Fig. 4F, we then calculated the fraction of activity patterns associated with symbol 1 in the different instances of ‘RRRL’ being classified as ‘R1’.

### Cross-progression analysis (Fig. 5)

Because the relative cluster rearrangements accompanying terminal cluster convergence (see Fig 6) begins as the animals start exiting the center port, analysis window for cross-projection experiments was limited from 250 msec before to 50 (rather than 250) msec after center port exit.

To quantify the observed alignment between ‘RRRL’ and ‘RRRLR’ neural graphs (Fig. 5B), we first obtained ordinal progression dimensions for the ‘RRRL’ component of the ‘RRRLR’ graph in the same manner as was done for Fig. 4B (see above). Its ability to report progression of ‘RRRL’ and ‘LLLR’ sequences was determined using rank-aligned version of the ROC classification above. Rank alignment was achieved by skipping step 1 (sorting of distributions along the classification dimension, agnostic of their rank order) and using the actual ordinal information to order distributions. Thus, two graphs in an anti-parallel configuration would show diminished rank-aligned cross-classification score.

### Analysis of terminal cluster ‘convergence’ (Fig. 6D)

Euclidean distances between cluster centroids – median location of all points in the cluster --were calculated in the corresponding PC spaces. Principal component analysis was done independently re-independently for each of the four analysis windows before the cluster conversion metric (distance ratio, see Fig. 6C) was calculated. Center out1 window was from 250 before to 50ms after center port exit. Center out 2 window was from 50 msec after to 350ms after around center port exit. Side in1 window was from 250 before to 50ms after side port entry. Side in2 window was from 50 after to 350ms after side port entry.

### Algorithmic interpretation of the ‘RRL+RRL’/ ‘LLR+LLR’ motif in forced pattern alternation (Fig. 7C)

Analyses were carried out in the high-dimensional PC space. First, centroid locations for the six clusters corresponding to individual symbols within ‘RRL+’ and ‘LLR+’ sequence instances were calculated. Subsequently, the closest of these six clusters was identified for the individual trials in all instanced of the second ‘RRL’ of the ‘RRL+RRL’ motif (or second ‘LLR’ in ‘LLR+LLR’ motif). Note that since the second RRL in ‘RRL+RRL-‘ (second ‘LLR’ in ‘LLR+LLR-‘) was never rewarded, its component symbols never contributed to the calculation of the six cluster centroids. For R1 and R2 of the second ‘RRL’ instances, we calculated the fraction of instances for which the closest cluster was “R1” and “R2” in from the three clusters of the ‘RRL+’ set respectively. For L3 of the RRL instances of interest, we calculated the fraction of instances for which the closest cluster was either “L3” of “RRL+”, or, separately, “L1” of “LLR+”.

---

**Fig. S1.**
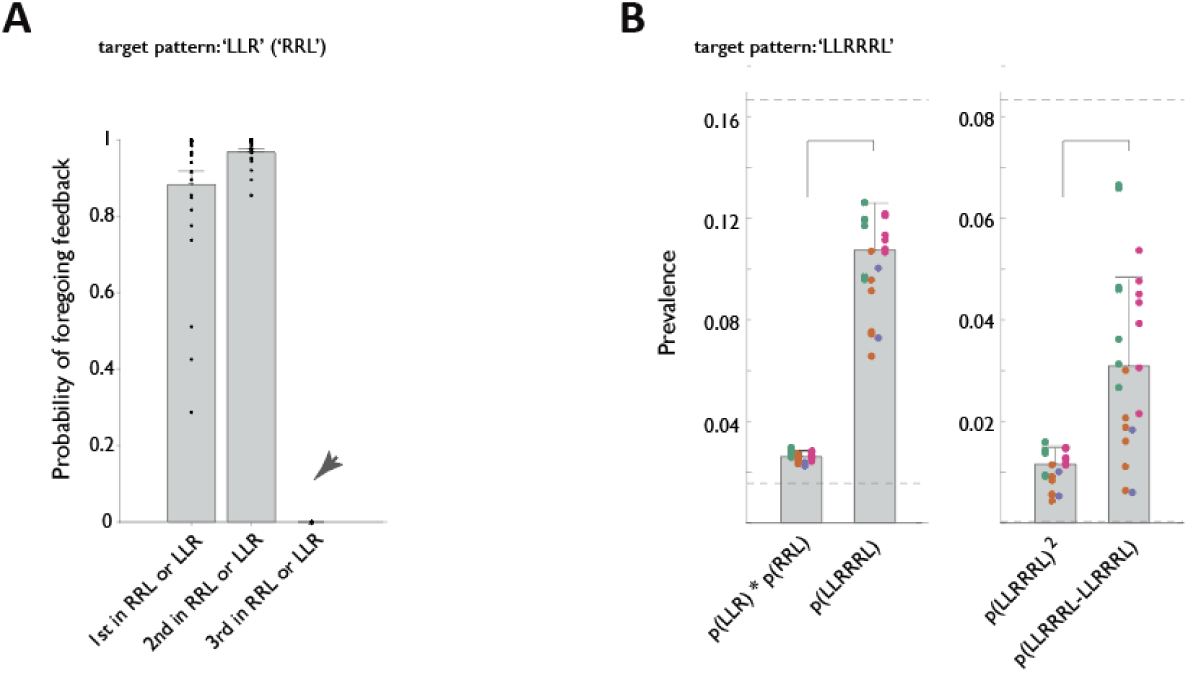
Feedback delays reveal flexible mental punctuation at sequence ends. (**A**) Probability that animals would forgo feedback on individual steps of minimal pattern matches in ‘LLR’ and ‘RRL’ contexts. Notice robust unwillingness to forgo delay on the terminal symbol (arrow). (**B**) Subsequent behavior of a subset of animals from (A) in the ‘LLRRRL’ latent context before the re-introduction of feedback delays in Fig. 1G,H. This stage was used to verify that animals had indeed discovered the concatenated latent target pattern, as indicated by the fact that the prevalence of ‘LLRRRL’ (or of its direct concatenation) greatly exceeds one that would be expected through a chance apposition of the ‘LLR’ and ‘RRL’ subcomponents.

**Fig. S2.**
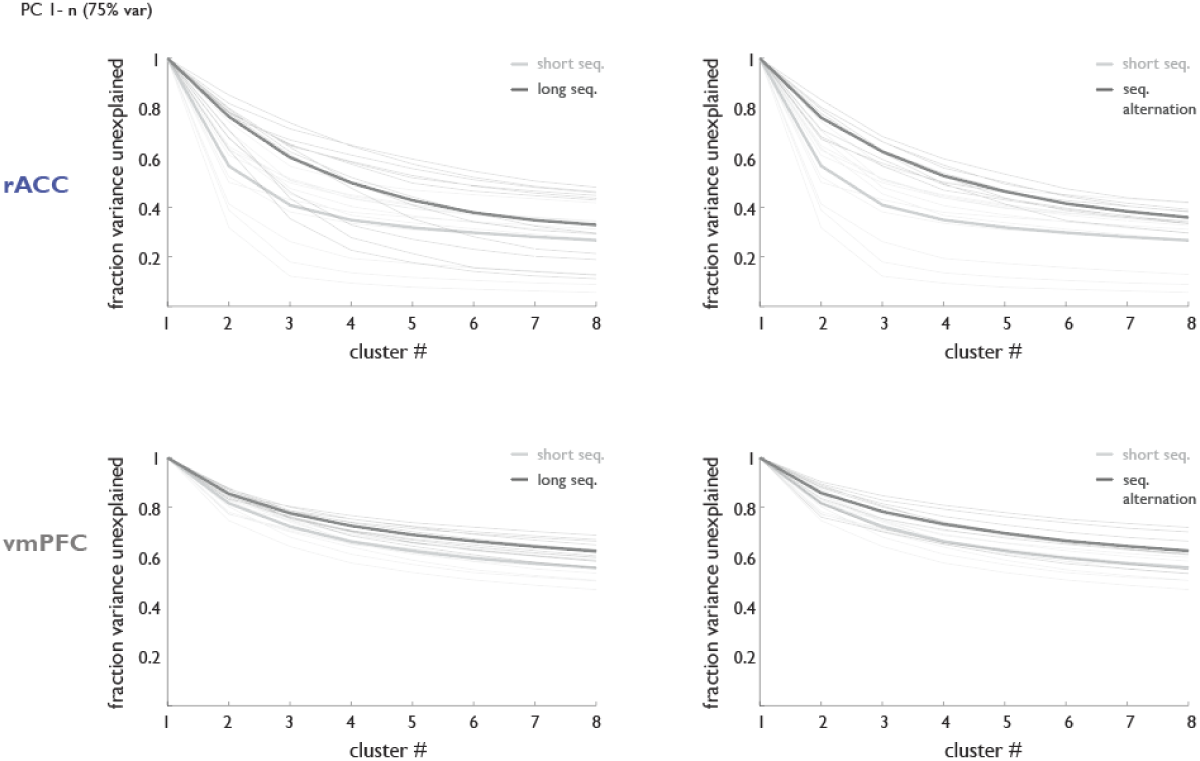
rACC but not vmPFC representation scales with task complexity. Same as in Fig. 2D, but in the high dimensional space. Fraction of rACC (top row) or vmPFC (bottom row) data variance left unexplained by a model with different cluster numbers for sessions with long vs short sequences as latent structure (left row, n=12 sessions, N-=4 animals), and session with blocks of a single short sequence or forced alternation between two short sequences as latent structures (n=9 session, N=3 animals). Note that sample sizes were matched for comparisons within the same session. Thin lines: individual sessions. Thick lines: session averages.

**Fig. S3.**
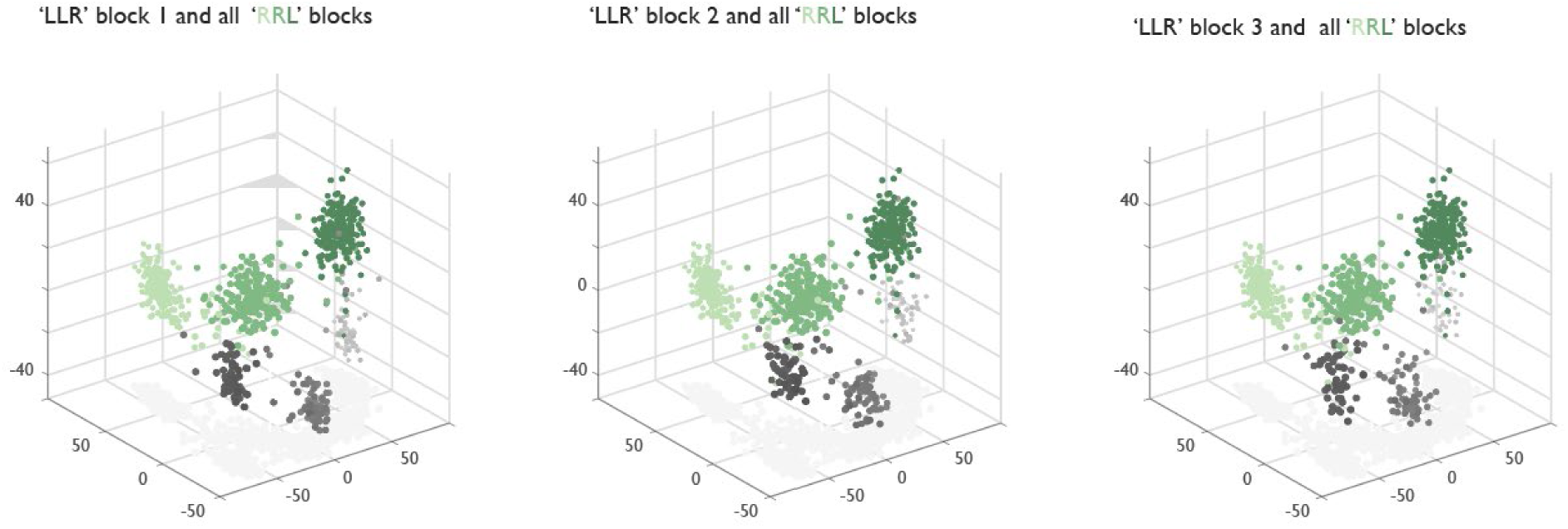
Instantiation of extra clusters is not related to elapsed time. Neural population geometry for three different ‘LLR’ blocks (in grey shading) and all three ‘RRL’ blocks from an example session with an interleaving ‘LLR’/’RRL’ block design.

**Fig. S4.**
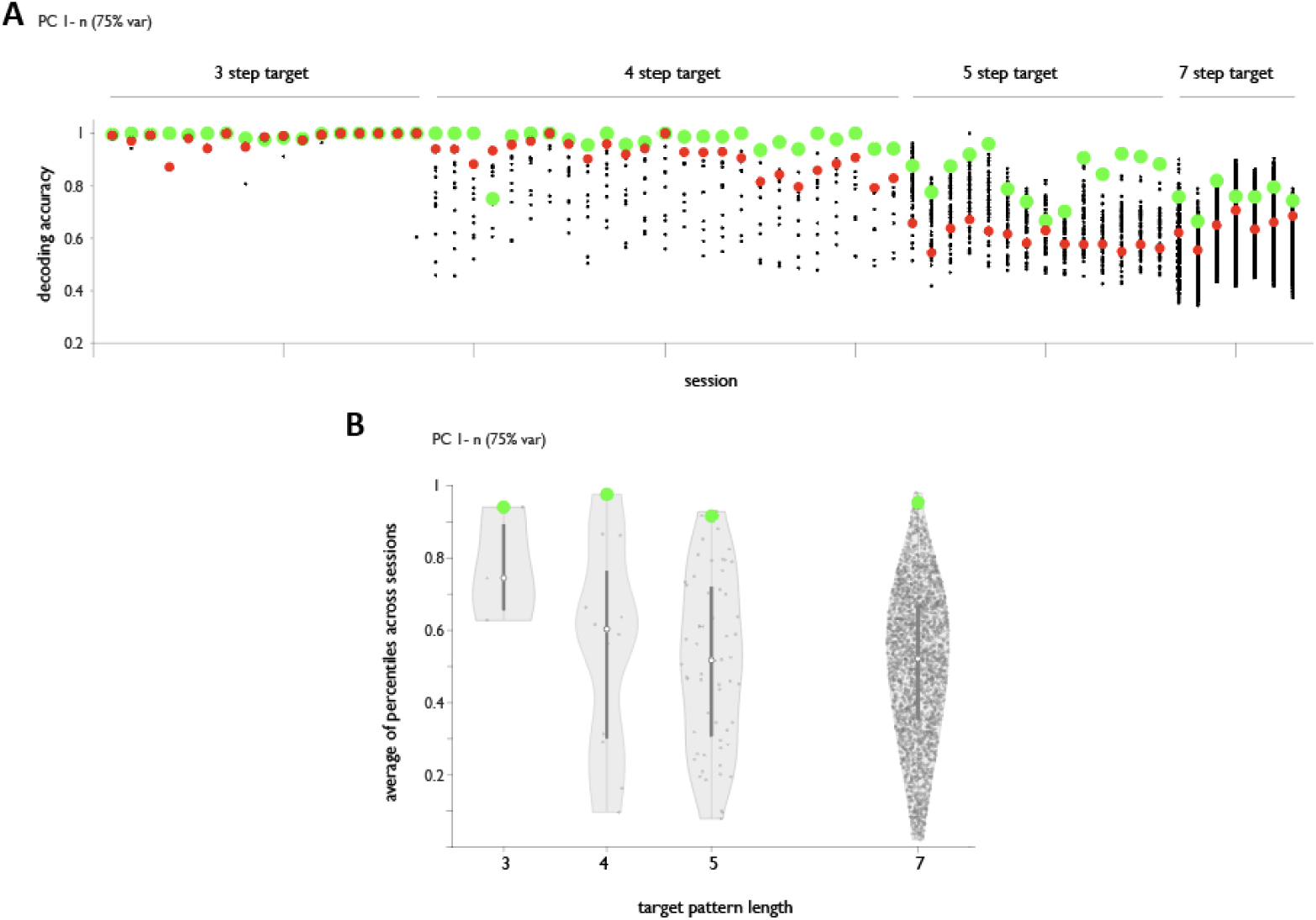
The ability to decode ordinal structure observed is not a trivial byproduct of easier separability in high-dimensional spaces. (**A**) Decoding accuracy for all possible ordinal re-indexing permutations in each behavioral session. Green: correct ordinal indexing, Red: Permutation with maxima index mixing. (**B**) Violin plot for the distribution of average percentile performances for all permutations. Green: correct ordinal indexing. [absolute rank of the correct indexing variant 1 out of 3, 1 out 12, 3 out of 60, 5 out of 2520].

**Fig. S5.**
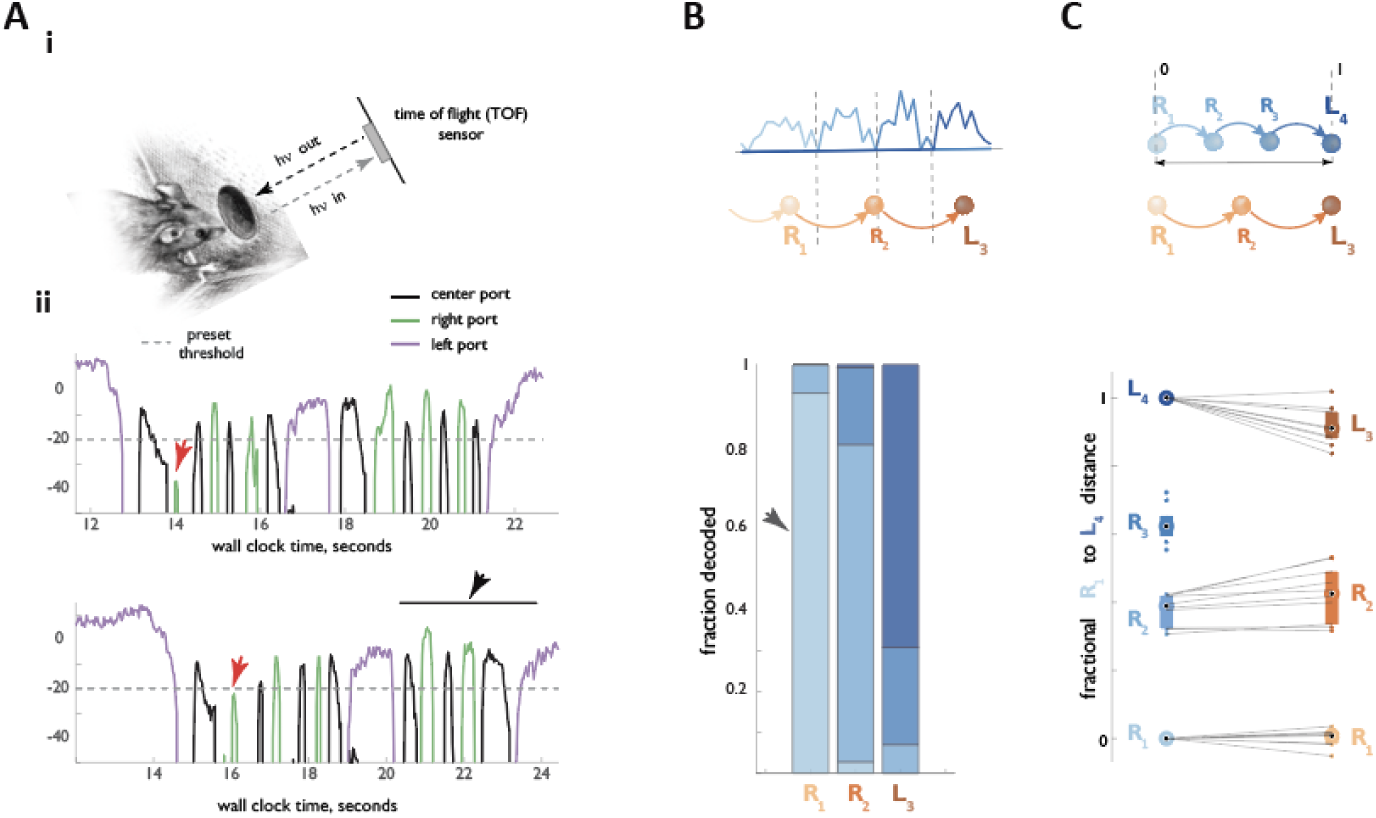
Restricting the putative ‘RRL’ dataset to registered ‘RRL’ instances NOT preceded by an unregistered ‘R’ reveals greater stability of the anchoring R1 node. (**A**) Schematic of unregistered behavioral symbol detection. (i) Detailed information on the depth of interaction with the nose port established with the help of a Time-of-flight (TOF) sensor. (ii) Two example TOF traces with legitimate (in their compound center port-side port structure) ‘R’s not registered by the hardware because animal’s interaction with the side port was not deep enough to cross the threshold (red arrowheads). Note also a legitimate ‘RRL’ in the second trace (black arrowhead). (**B**,**C**) Same as in Fig. 4J,K, but with the set of ‘RRL’s restricted to those that were not preceded by an unregistered ‘R’. Note the improved stability of the anchoring R1 node (arrowhead).

